# A pan-cancer atlas of somatic mutations in miRNA biogenesis genes

**DOI:** 10.1101/2020.07.22.216499

**Authors:** Paulina Galka-Marciniak, Martyna Olga Urbanek-Trzeciak, Paulina Maria Nawrocka, Piotr Kozlowski

## Abstract

It is a well-known and intensively studied phenomenon that the levels of many miRNAs are differentiated in cancer. miRNA biogenesis and functional expression are complex processes orchestrated by many proteins cumulatively called miRNA biogenesis proteins. To characterize cancer somatic mutations in the miRNA biogenesis genes and investigate their potential impact on the levels of miRNAs, we analyzed whole-exome sequencing datasets of over 10,000 cancer/normal sample pairs deposited within the TCGA repository. We identified and characterized over 3,600 somatic mutations in 29 miRNA biogenesis genes and showed that some of the genes are overmutated in specific cancers and/or have recurrent hotspot mutations (e.g., *SMAD4* in PAAD, COAD, and READ; *DICER1* in UCEC; *PRKRA* in OV; and *LIN28B* in SKCM). We identified a list of miRNAs whose level is affected by particular types of mutations in either *SMAD4, SMAD2*, or *DICER1* and showed that hotspot mutations in the RNase domains in DICER1 not only decrease the level of 5p-miRNAs but also increase the level of 3p-miRNAs, including many well-known cancer-related miRNAs. We also showed an association of the mutations with patient survival. Eventually, we created an atlas/compendium of miRNA biogenesis alterations providing a useful resource for different aspects of biomedical research.

## INTRODUCTION

Since the first reports of microRNA (miRNA) contributions to B-cell chronic lymphocytic leukemia [1], we have observed a substantial increase in reports describing the role of these small regulatory RNA molecules in different human diseases, including cancers (summarized in [2]). It was demonstrated that miRNAs are globally downregulated in cancer [3] and that upregulation or downregulation of certain miRNAs acting either as oncogenes (oncomiRs) or tumor suppressors (suppressormiRs) may contribute to cancer development and progression [4,5]. Numerous miRNA profiling studies have led to the identification of many miRNAs specifically altered in different types or subtypes of cancer. Many of these miRNAs play an important role in carcinogenesis and the regulation of different cancer-related processes, such as cell growth and differentiation, cell migration, apoptosis, and epithelial-to-mesenchymal transition (MET). Additionally, many miRNAs have been implicated as diagnostic and prognostic biomarkers and/or as potential therapeutic targets in cancer (e.g., [6–9]).

miRNAs are generated through a multistage process of miRNA biogenesis tightly controlled by various proteins consecutively nursing primary miRNA transcripts (pri-miRNAs) from their transcription to their cellular function within the miRNA-induced silencing complex (miRISC) [10–13]. The major steps of canonical miRNA biogenesis include nuclear pri-miRNA processing by the microprocessor complex, whose core is formed by RNase DROSHA acting together with DGCR8 and several other regulatory proteins, including P68 (DDX5) and P72 (DDX17), to release from the pri-miRNA hairpin-shaped secondary precursor (pre-miRNA). Next, pre-miRNA is exported to the cytoplasm by the Exportin-5(XPO5):Ran-GTP(RAN) complex, where it is intercepted by the multiprotein miRISC loading complex (RLC) containing the RNase DICER1, which cuts off the pre-miRNA apical loop to release an ~22-bp-long miRNA duplex. Within the miRISC, the miRNA duplex is unwound (supported by, i.a., GEMIN4 and MOV10) to select a miRNA guide strand (mature miRNA) that recognizes mRNA targets by complementary interaction and silences them with the assistance of AGO and TNRC6A (GW182) proteins by translation repression and/or RNA deadenylation and degradation. Each step of this process may be further regulated by additional mechanisms/proteins that either increase or decrease the miRNA biogenesis rate [11,14,15]. For example, LIN28A/B binds to the apical loop of specific pre-miRNAs, including pre-let-7, and upon its uridylation by ZCCHC11 or ZCCHC6 TUTases leads to pre-miRNA degradation by DIS3L2 exonuclease [16,17]. It should also be noted that there are alternative pathways of miRNA biogenesis, such as the generation of miRNAs (mirtrons) from specific short introns (DROSHA-independent) [18] or DICER1-independent processing of miR-451a by AGO2 [19,20].

There are currently more than 2,600 human miRNAs deposited in miRBase [21,22]. It is speculated that miRNAs regulate the expression of most protein-coding genes [23,24]. miRNA levels and consequently levels of controlled genes may be affected by various factors and processes. First, the expression of miRNA genes, such as the expression of protein-coding genes, is regulated by various transcription factors, such as MYC, TP53, or SMAD4 [25–28]. It was shown that many miRNA genes are located in copy number-variable (CNV) regions and are frequently amplified or deleted in cancer [29–31]. miRNA genes may also be affected by aberrant DNA methylation and histone acetylation, leading to the silencing of the miRNA genes [32–38].

Additionally, methylation of miRNA precursors may facilitate their processing [39] as well as impair the ability of miRNAs to downregulate their targets [40]. It was also shown that the occurrence of single nucleotide polymorphisms (SNPs) [41–44] and germline or somatic mutations [45–49] may affect miRNA processing (level) and the ability of miRNAs to recognize their targets. Mature miRNAs may be captured and inactivated by cellular miRNA sponges such as lncRNAs, circRNAs, or pseudogenes [50,51]. Finally, the deficiency or impairment of the components of miRNA gene transcription activators, miRNA processing machinery, and the miRISC complex (for simplicity, cumulatively called miRNA biogenesis genes/proteins) described in the previous paragraph may affect miRNA levels and the effectiveness of miRNA gene silencing, respectively [52,53].

A body of evidence has indicated that deleterious germline mutations in the *DICER1* gene are responsible for DICER1 syndrome, an inherited disorder characterized by an increased frequency of various types of malignant and benign tumors that occur predominantly in infants and young children, the most common and most characteristic of which is pleuropulmonary blastoma [54,55]. It was shown that in cancers associated with DICER1 syndrome as well as other early childhood cancers (e.g., Wilms’ tumor), a specific pattern of somatic *DICER1* second-hit missense mutations occurs. All these mutations are located in or adjacent to metal-ion-binding residues (hotspots; predominantly D1709 and E1813) of the RNase IIIb domain (RIIIb) [54]. Later, in similar types of childhood cancers, a similar pattern of somatic mutations was also identified in the corresponding residues (E1147, D1151) of the RIIIb in *DROSHA* [56–59]. Functional analyses revealed that the mutations in *DICER1* lead to less effective generation of 5p-miRNAs [60–62], whereas mutations in *DROSHA* affect the generation of miRNAs from both pre-miRNA arms [56], as reviewed in [54,63,64]. Very recently, it was also shown that the recurring mutation in the RNase IIIa domain (RIIIa) of *DICER1* occurring predominantly in uterine carcinoma may cause the same effect as the mutations in RIIIb [65]. Interestingly, the occurrence of *DROSHA* mutations in Wilms’ tumor coincides with the occurrence of mutations in *SIX1* and *SIX2*, transcription factor genes that are also frequently mutated in the tumor [66]. Another hotspot mutation commonly occurring in Wilms’ tumor is E518K in the double-stranded RNA-binding domain (dsRBD) of DGCR8 [57,58,66]. It was also shown that in cancers with a high rate of microsatellite instability (MSI), such as colon, gastric, and endometrial tumors, specific indel hotspots occur in *TRBP* and C-terminal positions of *XPO5* [67,68]; however, these mutations were not further analyzed in other studies. Knowledge of the germline and somatic variation in miRNA biogenesis genes is summarized in [64]. Additionally, *SMAD4*, encoding the SMAD4 transcription factor activating many genes in response to transforming growth factor beta (TGFB)/bone morphogenetic protein (BMP) signaling [69,70], is a well-known tumor suppressor gene that is highly mutated in many cancers, including pancreatic and colorectal cancers [71]. Although SMAD4 was also implicated in the transcription of miRNA genes [25,72,73], the effect of *SMAD4* mutations has never been tested in the context of the activation of miRNA genes. Furthermore, it was shown that some SNPs in miRNA biogenesis genes are associated with the risk of various cancers. Examples include (i) the rs3742330 (A>C) SNP located in the 3’ UTR of *DICER1* that affects DICER1 mRNA stability and is associated with susceptibility and malignancy in gastric cancer [74,75], increased survival of T-cell lymphoma patients [76] and lower prostate cancer aggressiveness [77]; (ii) the rs78393591 SNP in *DROSHA* and rs114101502 SNP in *ZCCHC11* (TUTase responsible for pre-miRNA uridylation and subsequent DICER1 cleavage inhibition) associated with the risk of breast cancer [78]; and (iii) the rs11786030 and rs2292779 SNPs in *AGO2*, rs9606250 SNP in *DGCR8*, and rs1057035 SNP in *DICER1* associated with the survival of breast cancer patients [79]. Additionally, the SNPs rs2740348 C>G and rs7813 C>T in *GEMIN4*, a gene involved in miRISC formation and miRNA-duplex unwinding, were implicated in the risk of several cancers, although the results were not conclusive [80–84].

In this study, we took advantage of the data generated within The Cancer Genome Atlas (TCGA) project to analyze the somatic mutations in miRNA biogenesis genes. As a result, in a wide panel of 33 cancer types consisting of over 10,000 samples, we identified hundreds of mutations and many recurrently mutated hotspot positions and showed that some of the genes are specifically overmutated in particular cancer types. We also confirmed the common occurrence of deleterious mutations in *SMAD4* and further characterized the specific hotspot mutations in *SMAD4, SMAD2*, and *DICER1*, the last group of which were previously reported mostly in childhood cancers. We followed up on the consequences of some of the mutations and showed characteristic changes in miRNA profiles resulting from specific mutation types in *DICER1, SMAD4*, and *SMAD2*. We also showed the associations of the mutations with cancer characteristics and patient survival. Additionally, the specific hotspot mutations in *DROSHA* and *DGCR8* commonly observed in Wilms’ tumor and other childhood cancers were absent in adult cancers.

## METHODS

### Data resources

We used molecular and clinical data (Level 2) for 33 cancer types generated and deposited in the TCGA repository (http://cancergenome.nih.gov). These data included the results of somatic mutation calls in whole-exome sequencing (WES) datasets of 10,369 samples (later limited to 10,255) analyzed against matched normal (noncancer) samples with the use of the standard TCGA pipeline. Copy number data were obtained via Xena UCSC as a ‘gene-level copy number (gistic2_thresholded)’ dataset of the TCGA Pan-Cancer (PANCAN) cohort. The crystal structure of the phosphorylated Smad2/Smad4 heterotrimeric complex (PDB code: 1U7V) [85] was visualized with the use of PyMOL (Schrödinger, LLC, New York, NY, USA).

### Data processing

We analyzed somatic mutations in 29 miRNA biogenesis genes (coding exons were extended by 2 nt on each side to enable identification of definitive splicing mutations). The genomic coordinates of the analyzed genes/regions are shown in Supplementary Table S1. From the WES data generated with the use of four different algorithms (MuSE, MuTect2, VarScan2, and SomaticSniper), we extracted somatic mutation calls with PASS annotation. The extraction was performed as described in [45] with a set of in-house Python scripts available at (https://github.com/martynaut/mirnaome_somatic_mutations). Briefly, the lists of somatic mutations detected by different algorithms were merged such that variants detected by more than one algorithm were not multiplicated. To further increase the reliability of the identified somatic mutations (and avoid the identification of uncertain mutations), we additionally removed those that did not fulfill the following criteria: (i) at least two alternative allele-supporting reads in a tumor sample (if no alternative allele-supporting read was detected in the corresponding normal sample); (ii) at least a 5x higher frequency of alternative allele-supporting reads in the tumor sample than in the corresponding normal sample; (iii) a somatic score parameter (SSC) > 30 (for VarScan2 and SomaticSniper); and (iv) a base quality (BQ) parameter for alternative allele-supporting reads in the tumor sample > 20 (for MuSE and MuTect2). All mutations were designated according to HGVS nomenclature at the transcript and protein levels, and the effects of mutations were predicted using the Ensembl Variant Effect Predictor (VEP) tool [86]. For visualization of mutations on the gene maps, we used ProteinPaint from St. Jude Children’s Research Hospital - PeCan Data Portal [87]. The protein domains visualized on gene maps were positioned according to UniProt [88]

### miRNA expression analysis

We obtained miRNA expression data via Xena UCSC as batch-effect normalized data for TCGA Pan-Cancer data (note that due to normalization, some miRNA levels were below 0). Expression data from the set of ~700 miRNAs were filtered to exclude miRNAs with undetectable signals (level=0) in more than 30% of pan-cancer samples (or 10% when the analysis was performed for specific cancers). Next, to enable pan-cancer comparisons, we normalized the variation (range of −1 to 1) and median (median=0) of miRNA levels to be equal in each cancer type. The normalized miRNA level changes (in pan-cancer) were calculated/expressed as differences, and the changes in raw (non-pan-cancer-normalized) miRNA levels (in individual cancers) were calculated/expressed as log2 fold changes.

### Statistics

Unless stated otherwise, all statistical analyses were performed with statistical functions in the Python module scipy.stats. Particular statistical tests are indicated in the text, and if not stated otherwise, p<0.05 was considered significant. If necessary, p-values were corrected for multiple tests. Mutation density was calculated as the number of detected mutations divided by the length of analyzed genes (total length of all coding exons). For patient survival analyses, we used a log-rank test (from the lifelines library [89]). To determine the direction of mutation effects on survival, we used Cox’s proportional hazard model. Survival plots were created using KaplanMeierFitter from the lifelines library. A comparison of the occurrence of mutations in miRNA biogenesis genes among cancer stages (with correction for cancer type) was performed with the Cochran-Mantel-Haenszel test for pan-cancer and Fisher’s exact test for specific cancers.

## RESULTS

### Distribution of somatic mutations in miRNA biogenesis genes across different cancer types

For analysis, we selected 29 miRNA biogenesis genes encoding proteins playing roles in (i) the transcription of primary miRNA precursors (pri-miRNAs), (ii) pri-miRNA to pre-miRNA processing in the nucleus, (iii) the export of pre-miRNA from the nucleus to the cytoplasm, (iv) pre-miRNA processing and miRNA maturation in the cytoplasm, and (v) miRNA:target recognition/interaction and regulation of downstream silencing effects (Table 1). Cumulatively, our panel included 465 protein-coding exons covering ~75 kbp (Supplementary Figure S1).

**Table 1.** List and characteristics of the selected miRNA biogenesis genes.

| Gene/Protein ID<br>(alias) | miRNA-related function |
| --- | --- |
| **Nuclear steps of miRNA biogenesis** |  |
| SMAD4 | Activates miRNA precursor transcription upon TGFB/BMP activation [120,124] |
| FUS | Facilitates cotranscriptional DROSHA recruitment to pri-miRNAs; shown to bind a terminal loop of specific neuronal pri-miRNAs and to enhance their processing [128] |
| SRSF3 (SRP20) | Enhances mammalian pri-miRNA processing upon binding to the CNNC motif in the 3p flanking sequence [129] |
| DROSHA | RNase III; catalyzes pri-miRNA to pre-miRNA for processing/cleavage [11,130,131] |
| DGCR8 | Cofactor of DROSHA; coordinates the recognition of pri-miRNA at the dsRNA-ssRNA junction; functions as a molecular anchor and direct DROSHA to cleave pri-miRNA 11 bp before the junction [11,132] |
| DDX5 (P68) | Plays a role in recognition/binding of pri-miRNAs by the DROSHA complex; recruits DROSHA/DGCR8 to pri-miRNA [133,134] |
| DDX17 (P72) | Plays a role in recognition/binding of pri-miRNAs by the DROSHA complex [133,135] |
| GSK3B | Facilitates pri-miRNA binding by DROSHA and enhances DROSHA association with cofactors DGCR8 and P72 [136]; DROSHA phosphorylation/stabilization [137] |
| SMAD2 | Accelerates pri-miRNA processing by DROSHA; in complex with SMAD4 activates miRNA precursor transcription upon TGFB/BMP signaling [72,124,125] |
| **Export to cytoplasm** |  |
| XPO5 (EXP5) | Plays a role in the nuclear export of pre-miRNA to the cytoplasm [138,139]; facilitates the nuclear cleavage of clustered pri-miRNAs [140] |
| RAN | Interacts with XPO5; plays a role in the export of pre-miRNA to the cytoplasm [138] |
| **Cytoplasmic steps of miRNA biogenesis** |  |
| DICER1 (DICER) | RNase III; catalyzes pre-miRNA to miRNA-duplex processing by cutting off the pre-miRNA terminal loop [11,141–143] |
| TARBP2 (TRBP) | Coordinates pre-miRNA recognition by DICER1 and the precision of DICER1 cleavage [144,145] |
| PRKRA (PACT) | Coordinates pre-miRNA cleavage by DICER1 and assures the precision of the cleavage; plays a role in miRISC assembly and thus participates in miRNA stabilization and accumulation in the cell [145,146] |
| ADAR | Double-stranded RNA-specific adenosine deaminase; plays a role in pri- and pre-miRNA stem editing (A to I), which makes miRNA precursors resistant to DICER1 cleavage [147] |
| KHSRP (FUBP2, KSRP) | Binds to the terminal loop sequence of a subset of miRNA precursors, promoting their maturation [148] |
| LIN28A and LIN28B | Bind to the terminal loops of specific pre-miRNAs (including pre-let-7 and pre-miR-9) and, upon recruitment of ZCCHC11 or ZCCHC6 that induces pre-miRNA uridylation, inhibit DICER1 processing [17,149,150] |
| ZCCHC11 (TUT4) and ZCCHC6 (TUT7) | Play a role in pre-miRNA uridylation and thus inhibit DICER1 processing [17,151] |
| DIS3L2 | Exoribonuclease; targets the uridylated let-7 precursors [16] |
| **miRNA functioning** |  |
| AGO1, AGO2, AGO3, and AGO4 | Play a role in miRISC formation/loading (summarized in [152,153]); catalytically active AGO2 functions as an endonuclease upon complementary mRNA:miRNA interaction [154] |
| GEMIN4 | Binds to the miRNA guide strand and facilitates the formation of a miRISC by unwinding the miRNA duplex [82,155] |
| MOV10 | Upon interaction with the miRNA-loaded AGO-protein complexes, plays a role in mRNA degradation; present in p-bodies [156,157] |
| FMR1 | Interacts with DICER1 and AGO1 during mRNA degradation [158]; controls DROSHA expression [159] |
| TNRC6A (GW182) | Component of P-bodies; plays a role in mRNA degradation upon interaction with Argonaute proteins [160,161] |

To identify somatic mutations in the miRNA biogenesis genes, we took advantage of WES datasets of 10,369 paired tumor/normal samples generated within the TCGA project. The collected samples cover 33 different cancer types, including 10 rare cancers (analyzed together as a pan-cancer cohort) (Supplementary Table S2). The list of all cancer types (full names and abbreviations) is shown in Figure 1 (to avoid confusion, we will use the abbreviations for the TCGA sample sets but not generally for particular types of cancer; in the latter case, we will use full cancer-type names). Applying the rigorous criteria described in the Materials and Methods, we identified a total of 5,483 mutations in the pan-cancer cohort. However, a substantial fraction (n=1834, ~30%) of the mutations were identified in a relatively small number (n=114, ~1%) of hypermutated samples, defined as samples with >10,000 mutations in the whole exome. As shown in Supplementary Figure S2, the number of mutations in hypermutated samples strongly depends on the general burden of mutations in these samples, implying enrichment of random, most likely passenger mutations in the hypermutated samples. Therefore, to reduce the proportion of confounding mutations, we removed hypermutated samples from subsequent analyses. The removed, hypermutated samples originated mostly from SKCM, UCEC, and COAD (Supplementary Figure S2).

**Figure 1.**
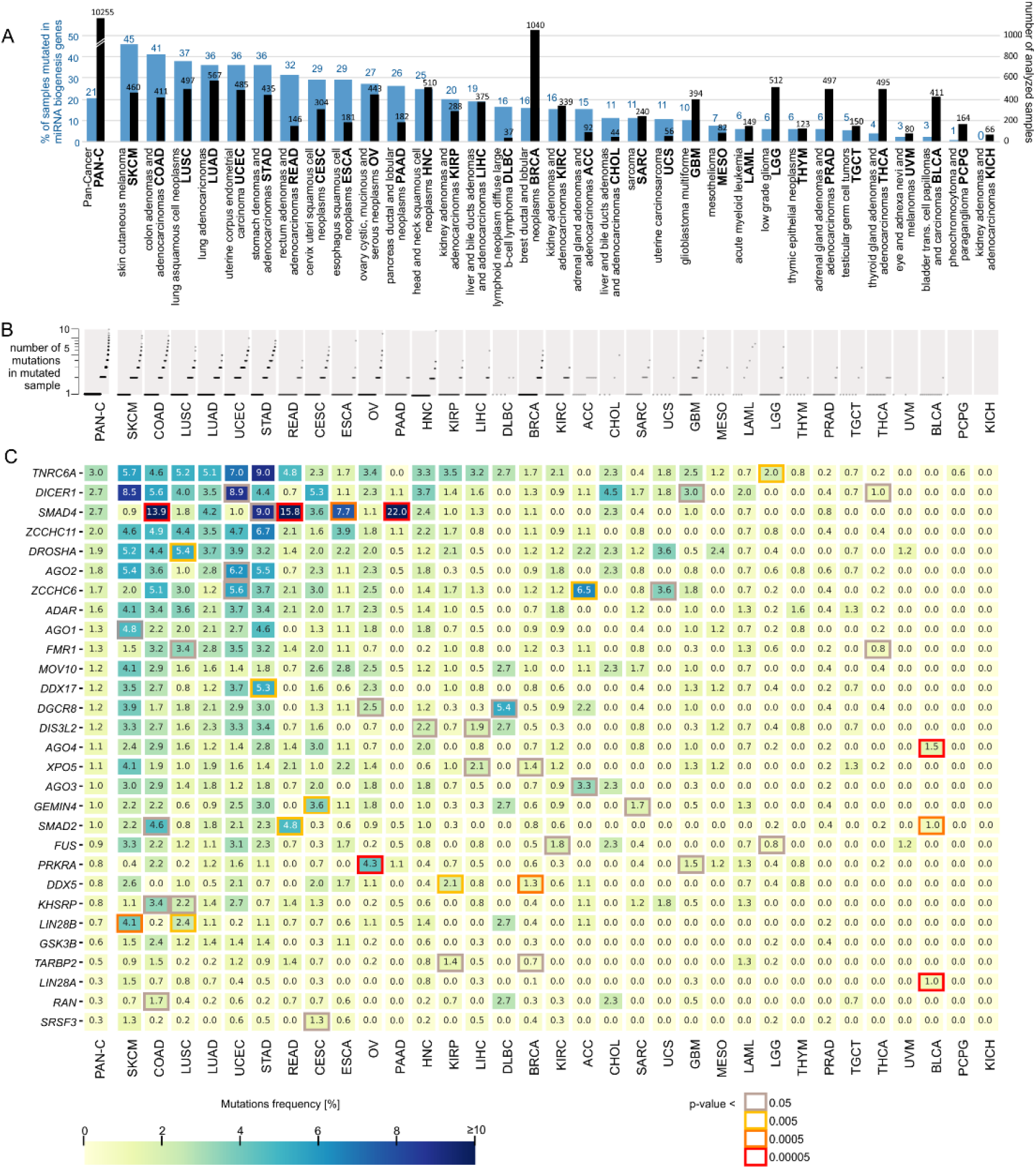
Mutation frequency in miRNA biogenesis genes across the analyzed cancer types. A) The total number of samples (black bars) and percentage of samples with mutations (blue bars) in the panel of miRNA biogenesis genes. B) The proportion of samples with different numbers of mutations. Each sample is shown as a dot (only samples with at least one mutation are shown). Due to the large number of samples with a particular number of mutations, dots overlap with each other. C) Heatmap showing the frequency [%] of mutations within each of the miRNA biogenesis genes (y-axis) in different cancer types (x-axis). The genes are ordered by the frequency of mutations in pan-cancer (first column). The genes significantly overmutated in particular cancer types are marked with a color frame indicating the nominal p-value (Fisher’s exact test; the p-value scale is indicated under the heatmap). A p-value<0.00005 (red frame) corresponds to significant enrichment after correction for multiple tests (adjusted p<0.05). Specific p-values are shown in Supplementary Table S4.

After removing the hypermutated samples, we continued analysis on 10,255 samples with 3,649 mutations, including 2,196 (60%) missense mutations, 774 (21%) synonymous mutations, and 625 (17%) definitive deleterious mutations, consisting of 341 frameshift, 222 nonsense, and 62 splice-site mutations (Supplementary Table S3). Other types of mutations, such as start or stop codon mutations or complex mutations, were also present in small fractions. At least one mutation was detected in 2,104 (21%) samples, including numerous samples with more than one mutation (Figure 1A and 1B). The frequency of mutated samples differed substantially among cancer types, ranging from 45% (SKCM) to 0% (KICH) (Figure 1A, Supplementary Table S3), and roughly corresponded to the mutational burden in particular cancer types.

### Frequency of somatic mutations in miRNA biogenesis genes across 33 cancers

As shown in Figure 1C, *TNRC6A, DICER1*, and *SMAD4* are among the most highly mutated genes (~3% of samples in pan-cancer), and the least mutated genes (~0.3%) are *LIN28A, RAN*, and *SRSF3*. Although there is some correlation between the frequency of mutations in the particular genes and the length of their protein-coding sequences (R^2^=0.62), the overall mutation frequency cannot be simply explained by the length of the genes. Additionally, none of the genes are well-known highly mutated genes or are located in late-replicating regions known to be overmutated in cancer [90].

Although there is a general correlation of mutation frequency in particular genes between cancers, there are also apparent striking exceptions of genes specifically overmutated in particular cancers (Figure 1C). This contrasts with observations that over- and undermutated regions generally overlap between different cancer types [90] and suggests the nonrandom occurrence and potentially functional nature of the overmutations. To identify overmutated genes, we statistically compared the frequency of mutations (overmutation) in particular genes (versus all other genes) in particular cancers with corresponding frequencies in pan-cancer (Figure 1C, Supplementary Table S4). The most striking example of an overmutated gene is *SMAD4* overmutated in PAAD (22% of samples), READ (16%), COAD (14%), STAD (9%), and ESCA (8%). Other interesting examples included *AGO4, LIN28A*, and *SMAD2* overmutated in BLCA (an otherwise extremely low-mutation cancer with no mutations in other genes); *LIN28B* overmutated in SKCM; *PRKRA* overmutated in OV; and *DDX5* overmutated in BRCA and KIRP. There are also other genes with increased mutation frequency, e.g., *TNRC6A* in STAD and *DICER1* in SKCM and UCEC (nominal p>0.005), but these overmutations are only nominally significant. Consistent with the above findings, the overmutated genes are outliers in terms of the correlation of mutation frequencies in particular genes between particular cancer types and the remaining pan-cancer (Supplementary Figure S3).

### Distribution of mutations in the miRNA biogenesis genes – identification of hotspot mutations

To illustrate the mutation distribution along the protein sequences, all the mutations were annotated according to HGVS nomenclature and visualized in lollipop plots (Figure 2 and Supplementary Figure S4). As shown in the plots, although most of the mutations are quite evenly distributed along the genes, there are also some hotspot regions/mutations suggesting the functional nature of these changes. The most striking example is a cluster of 8 hotspots of recurrently mutated amino acid (AA) residues (i.e., D351, G352, D355, P356, R361, H382, G386, and D537) occurring in the MH2 domain of SMAD4 (Figure 2A). The most prominent hotspot position is R361, which by itself acquired 37 missense mutations, accounting for 23% of all *SMAD4* missense mutations. There are also two recurrently mutated AA residues (i.e., P305 and R321) in the MH2 domain of SMAD2 (Figure 2B). The hotspot mutations in the MH2 domains (both in SMAD4 and SMAD2) likely affect SMAD4:SMAD2 heterotrimer formation. Additionally, there are two recurrent protein-truncating nonsense mutations in the MH2 domain of SMAD2, i.e., p.S306Ter and p.S464Ter, the latter of which occurs 13 times and truncates the protein just before two phosphorylation sites (S465 and S467) critical for activation of SMAD2 upon TGFB/BMP signaling [91].

**Figure 2.**
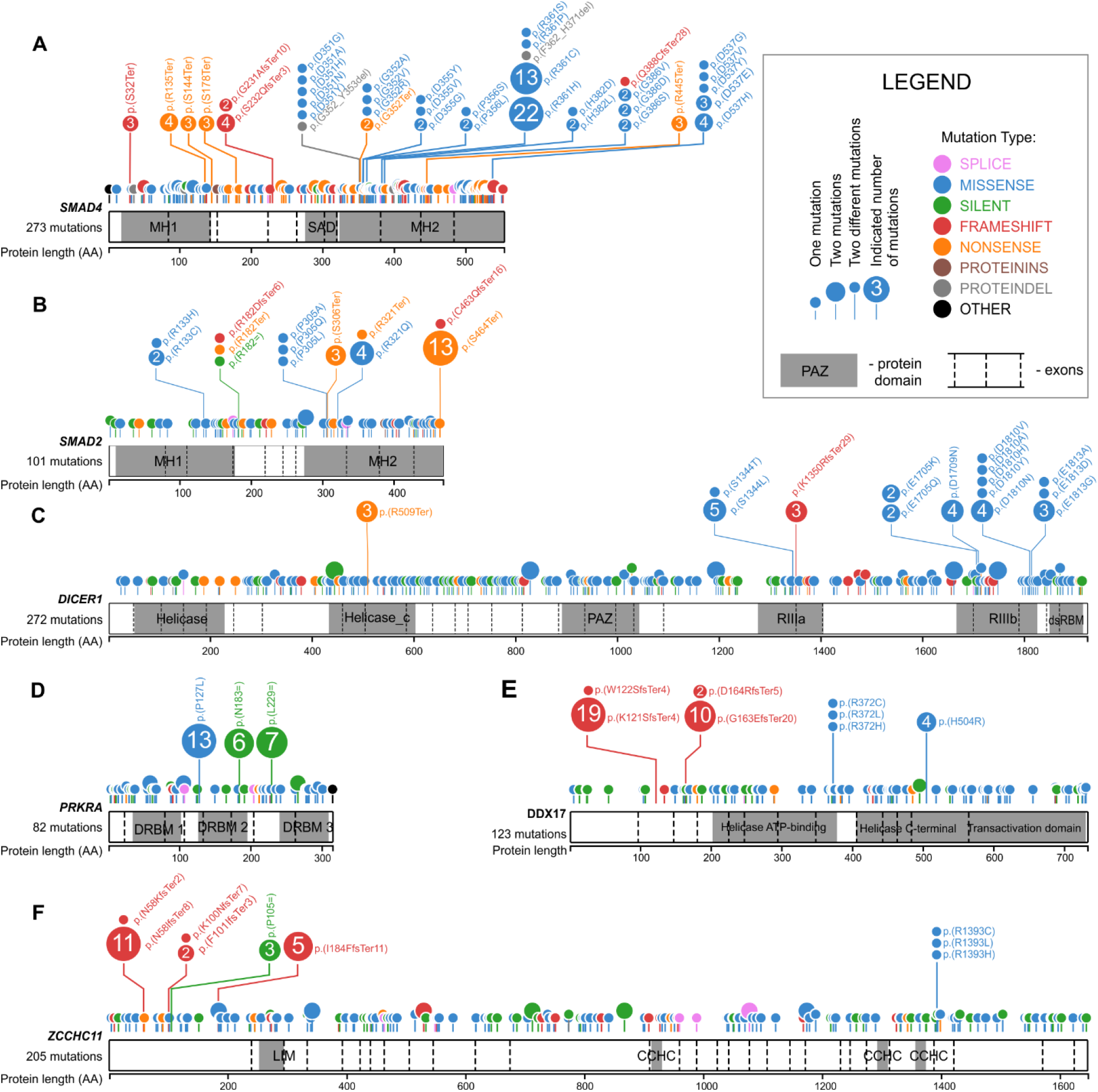
Distribution of the identified mutations in the miRNA biogenesis genes. **A)**, **B)**, **C)**, **D)**, **E)**, and **F)** depict *SMAD4, SMAD2, DICER1, PRKRA, DOX17*, and *ZCCHC11*, respectively. The remaining miRNA biogenesis genes are shown in Supplementary Figure S4. Mutations are visualized in the form of lollipop plots along the gene coding sequences, with the exon structure and protein functional domains indicated. The size of a mutation symbol (circle) is proportional to the number of mutations, and the color indicates the type of mutation (as shown in the legend). All mutations were annotated according to HGVS nomenclature, and the effect of the mutations at the protein level was denoted with the VEP tool (Ensembl).

Two other clusters of missense hotspot mutations are located in metal ion-binding residues of the RIIIa (S1344) and RIIIb (E1705, D1709, D1810, and E1813) domains of *DICER1* (Figure 2C). These hotspots were previously detected and functionally characterized in various pediatric cancers [54,64], thyroid adenomas [92], and the TCGA cohort, mostly in UCEC samples [65].

It is also worth noting the hotspot missense mutation, i.e., p.P127L in the DRBM 2 domain of *PRKRA* (Figure 2D), occurring 13 times in OV, COAD, GBM, and LUAD (6, 3, 3, and 1 mutations, respectively). As the DRBM 2 domain plays a role in interaction with other proteins (e.g., protein kinase R, PKR), the mutation may affect the interactions. Although the mutation overlaps with the SNP (rs75862065), the fact that it occurs predominantly in OV, which is otherwise a moderately mutated cancer, combined with the overall overmutation in PRKRA, argues against the accidental occurrences of the mutation as artifacts of the mutation calling process. There are also two hotspot synonymous mutations in FUS (p.G222= and p.G227=) (Supplementary Figure S4). A role of such hotspots cannot be excluded; however, we did not investigate them further in this study. Similarly, recurring missense mutations in the other genes may be important for specific cancers; however, they occur much less frequently.

There are also some hotspot indel mutations, e.g., p.N58IfsTer8 in *ZCCHC11*, p.Y948MfsTer16 in *ZCCHC6*, p.K121SfsTer8 and p.G163EfsTer20 in *DDX17*, p.W804GfsTer99 and p.R1183GfsTer7 in *TNRC6A*, p.L17CfsTer99 in *AGO1*, and pA603RfsTer71 in *AGO2* (Figure 2 and Supplementary Figure S4). However, it must be noted that the exact position of indel hotspot mutations may not be necessarily driven by cancer advantage but by sequence properties (e.g., the presence of short tandem repeat motifs). In fact, many indel mutations and some of the indel hotspots occur at sequence motifs often mutated as a result of MSI. Additionally, as most indels in coding sequences result in frameshifts and premature termination of translation, triggering nonsense-mediated mRNA decay (NMD) and leading to the complete loss of mRNA, the exact position of indel hotspots may not be that important.

In the next step, we looked for hotspot mutations previously observed in the miRNA biogenesis genes, i.e., (i) E969 and E993 in RIIIa and E1147, D1151, Q1187, and E1222 in RIIIb in *DROSHA*, (ii) p.E518K in the dsRDB1 domain in *DGCR8*, and (iii) p.R440Ter in *XPO5* observed in different pediatric cancers, especially in Wilms’ tumor [56,57,59]. Of these mutations, we found p.E518K in *DGCR8* occurring in two of 495 (0.4%) cases of THCA, p.R440Ter in *XPO5* in one case of UCEC and one case of SKCM, and p.D1151E in DROSHA in one case of COAD (Supplementary Figure S4). Of the indel hotspots in *XPO5* and *TARBP2* detected previously in colon, gastric, and endometrial tumors with MSI [67,68], we detected only two cases of the C insertion in the poly-C track (p.M145HfsTer13) in *TARBP2:* one in UCEC and one in STAD (both cancers often characterized by MSI).

### Functional consequences of *SMAD4 and SMAD2* mutations

From the visual investigation (Figure 1C and Figure 2A), it is apparent that *SMAD4* is the gene with the highest density of mutations (273 mutations, 160 mut/kbp) and the largest proportion of deleterious mutations (33%). As mentioned in the previous paragraph, there are also 8 hotspots of missense mutations, 7 of which are located in a relatively small region (D351 to G386) of the MH2 domain, playing a role in heterotrimer formation with other receptor-dependent SMAD proteins (R-SMADs, e.g., SMAD2) to mediate the TGFB/BMP transcriptional response [93,94]. The collection of a relatively large number of mutations associated with different cancer types reveals that the proportion of hotspot and deleterious mutations [in pan-cancer, 71 (44%) vs. 91 (56%), respectively] differs substantially between cancers (Figure 3A) and is the highest in READ (86% vs. 14%; p=0.001) and the lowest in PAAD (27% vs. 73%; p=0.032). This may suggest a different role of SMAD4 in different cancers and different effects of these two types of mutations. Most of the hotspots (5 of 8 hotspots, 83% of all hotspot mutations) coincide with charged (basic or acidic) AAs, affecting the electrostatic properties of MH2 important for interaction with R-SMADs [85]. To visualize the location of the hotspot AA residues, we marked them on a crystal structure of the heterotrimeric SMAD4:(SMAD2)2 complex [85]. As shown in Figure 3B, all hotspot residues are located on the surfaces of the SMAD4:SMAD2 interaction, which is consistent with the notion that hotspot mutations prevent SMAD complex formation and thus avert transcription of SMAD-controlled genes in response to TGFB/BMP signaling.

**Figure 3.**
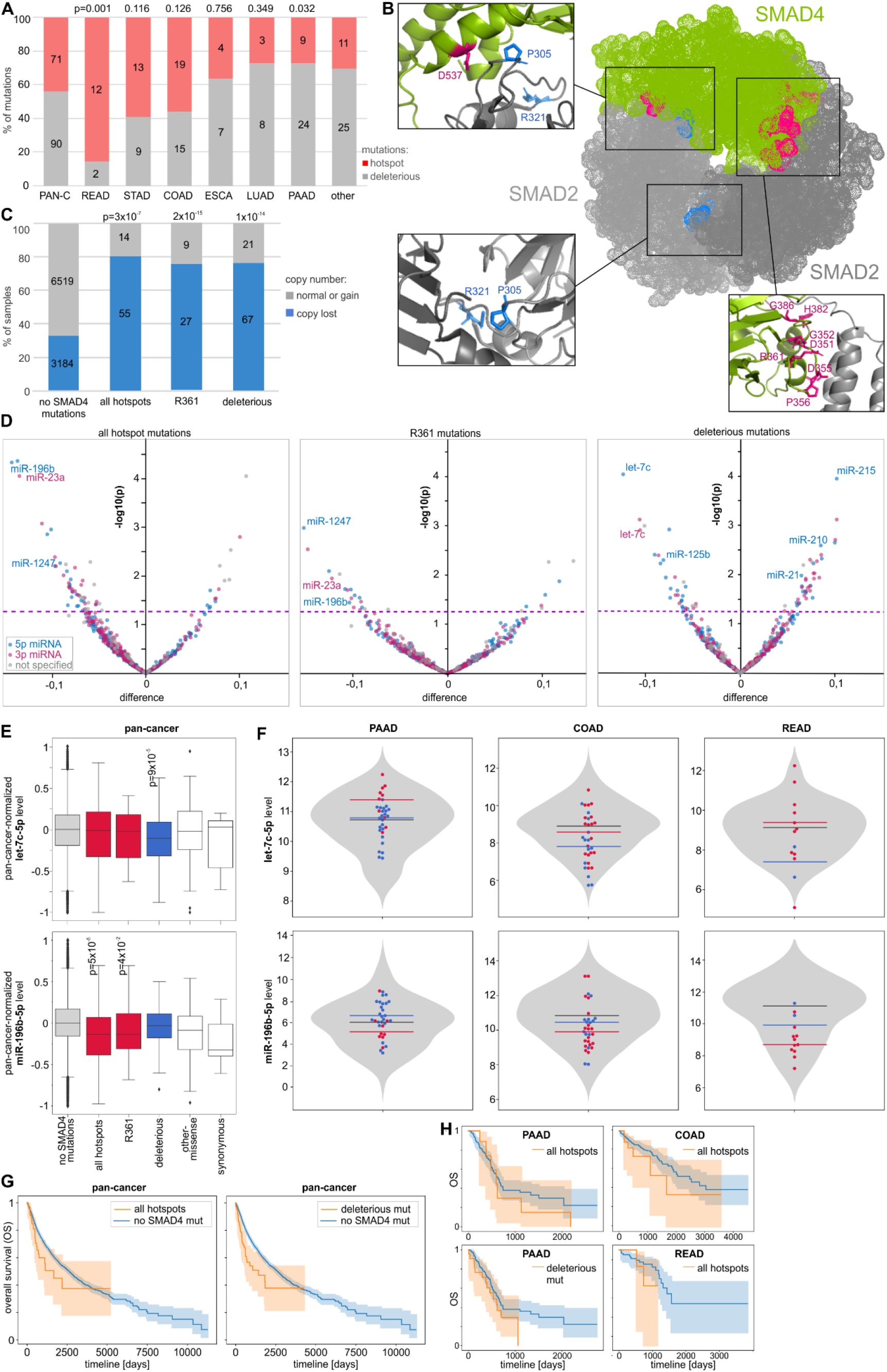
Characteristics of *SMAD4* mutations. A) The proportions of hotspot and deleterious mutations (y-axis) in different cancer types (x-axis). B) Localization of hotspot AA residues in the SMAD4:(SMAD2)2 heterotrimeric complex. *SMAD4* mutations are indicated in pink, whereas *SMAD2* mutations are indicated in blue. C) The proportion of copy number alterations of *SMAD4* (y-axis) in samples with different types of *SMAD4* mutations. D) Volcano plots depicting miRNA level alterations in samples with different types of *SMAD4* mutations (indicated above the graphs) compared to samples without any *SMAD4* mutations. Positive and negative values on the x-axis indicate increased and decreased miRNA levels in mutated samples, respectively. The y-axis indicates the p-value on a log10 scale, and the p=0.05 threshold is indicated by a pink dashed line. The y-axis indicates the log10 p-values. Blue, pink, and gray dots indicate 5p-miRNAs, 3p-miRNAs, and miRNAs whose arm was not specified, respectively. The IDs of selected miRNAs are indicated on the graphs. E) Boxplots showing the distribution of the selected miRNA levels (y-axes) in samples with different types of mutations vs. samples without any *SMAD4* mutations (x-axes). The p-values<0.05 are indicated on the graphs. F) Violin plots showing the distribution of non-pan-cancer-normalized levels (y-axis) of the miRNAs shown in E in samples without *SMAD4* mutations and specific samples with different types of *SMAD4* mutations (dots of different colors). Horizontal lines indicate median miRNA levels in samples with different types of mutations. G) and H) Kaplan-Meier plots indicating the OS of patients without and with a specified type of *SMAD4* mutation in pan-cancer and specific cancer types, respectively.

The copy number analysis of *SMAD4* showed that *SMAD4* deletions are significantly more frequent in samples with *SMAD4* mutations than in samples without such mutations, indicating frequent loss of the second allele [loss of heterozygosity (LOH) effect] in samples with both hotspot and deleterious mutations (Figure 3C). *SMAD4* mutations were previously shown to affect the expression of many genes. Although it was proposed that the TGFB/SMAD pathway also controls the expression of miRNAs [25,27,28,73,95,96], the effect of the mutations on miRNA levels has never been tested. To do so at the pan-cancer scale, we took into account only 522 miRNAs whose level was >0 in at least 70% of tested TCGA samples. We normalized miRNA levels to make the median (equal to 0) and variance of these levels comparable between cancers.. As shown in Figure 3D, there is an excess of downregulated miRNAs in samples with hotspot mutations (e.g., at the level of p-value<0.05, 78% and 22% of miRNAs are downregulated and upregulated, respectively), which is consistent with the expected impairment of SMAD complex formation and transcription factor activity. A similar effect is observed when the analysis is performed only for R361, the most frequently mutated hotspot residue (Figure 3D). Among the downregulated miRNAs (Supplementary Table S5), there are many miRNAs with well-documented cancer-related functions, for example, the recently discovered suppressormiRs miR-23a-3p (reviewed in [97]), miR-196b-5p [98,99], and miR-1247-5p [100], whose downregulation was associated with poor prognosis in breast cancer [101]. Noteworthy, such an effect of excessive miRNA downregulation is not visible for deleterious mutations. Among the most significantly downregulated or upregulated miRNAs in samples with deleterious mutations are let-7c-5p and miR-125b-5p or miR-215-5p, miR-21-5p, and miR-210-5p, respectively, all well-known cancer-related miRNAs (e.g., [102]). The frequent observation of pairs of 5p and 3p miRNAs (e.g., let-7c-5p and let-7c-3p) and groups of miRNAs generated from miRNA clusters (e.g., miR-99a/let-7c or miR-1/133a cluster) among the consistently altered miRNAs in samples with both hotspot and deleterious mutations indicates that at least some of the miRNA alterations result from aberrations at the transcriptional level. As shown in Figure 3E and 3F, the levels of let-7c-5p and miR-196-5p, which are examples of miRNAs most significantly altered in pan-cancer, often show similar alterations in the most commonly mutated cancers, i.e., PAAD, COAD, and READ (Figure 3E and 3F), although it must be noted that due to the very limited number of mutations, the results in individual cancers are not perfectly consistent and have to be interpreted with caution. To identify pathways/processes enriched in the genes regulated by the downregulated miRNAs, we performed KEGG pathway enrichment analysis with miRPath v3.0. The analysis showed that miRNAs downregulated both by the hotspot and deleterious *SMAD4* mutations are associated (adjusted p<0.01) with similar cancer-related processes, including ‘Pathways in cancer’, ‘ErbB signaling pathway’, ‘Glioma’, ‘TGF-beta signaling pathway’ and ‘Proteoglycans in cancer’ (Supplementary Table S6).

A comparison of *SMAD4* mutations with clinical cancer data did not show an association of mutations with tumor staging but showed a borderline significant association with decreased overall survival (OS) at the pan-cancer level (log-rank test: p=0.049 and p=0.0006 for hotspot and deleterious mutations, respectively) (Figure 3G). As the analyses performed at the pan-cancer level may be affected by the unequal distribution of mutations among cancer types, we repeated the survival analysis with the most frequently mutated individual cancers. Although the analyses of individual cancer types were of very limited statistical power because of a low number of mutations, some cancer types also showed trends toward decreased survival of patients with the mutations, i.e., COAD and READ (Figure 3H).

Although *SMAD2*, with a total of 101 mutations, including 35 deleterious mutations, is much less densely mutated (70 mut/kbp) than *SMAD4* (Figure 2B), it also contains recurrently mutated hotspot AA residues in the MH2 domain. The hotspots include two AA residues, i.e., P305 and R321 mutated 3 and 4 times, respectively, and the nonsense mutation p.S464Ter truncating the protein by 5 AAs, which is the most common *SMAD2* mutation (13 occurrences in cancers such as COAD, STAD, and BRCA). As in SMAD4, the SMAD2 hotspot residues localize at the SMAD4:SMAD2 interaction surfaces (Figure 3B).

We separately analyzed miRNA level profiles in samples with hotspot missense mutations (N=7), the p.S464Ter mutation (N=13), and deleterious mutations (not including p.S464Ter; N=22). As in samples with *SMAD4* mutations, samples with p.S464Ter and deleterious mutations showed a substantial excess of downregulated miRNAs [95% (34 of 36 altered miRNAs at p<0.05) and 70% (16 of 23), respectively]) (Figure 4A and Supplementary Table S5). The excess of downregulations is not visible for the hotspot missense mutations; however, it must be noted that due to a very low number of hotspot mutations, the analysis of miRNA levels for this type of mutation is of very low statistical power. Nonetheless, we observed one miRNA, miR-329-3p, playing a role in different cancers [103–106]), which was consequently downregulated in all three mutational groups. Other downregulated miRNAs playing a role in cancer include (i) miR-1247-5p and miR-1247-3p (in samples with p.S464Ter), recently identified as suppressormiRs [100]; (ii) miR-7-5p (in samples with deleterious mutations), a well-recognized suppressormiR, playing a role in downregulation of the growth, metastasis, and prognosis of various tumors (reviewed in [107]); and (iii) miR-380-5p (in samples with hotspot missense mutations), downregulating TP53 to control cellular survival [108]. As shown in Figure 4B and 4C, the examples of miR-329-3p and miR-380-5p illustrate that miRNA level changes identified in the pan-cancer analysis are also reflected in the most frequently mutated cancers, i.e., COAD and READ. SMAD2, similar to other R-SMADs, may also posttranscriptionally increase the level of a specific group of miRNAs, facilitating the processing of their pri-miRNAs by DROSHA [72]. Among the 44 miRNAs shown to be upregulated by R-SMADs or containing specific R-SMAD-interacting sequence motifs (23 of them tested in this study), miR-421 (p=0.01), miR-188-5p (p=0.03), and miR-877-5p (p=0.04) were downregulated in samples with the p.S464Ter mutation. A comparison of the SMAD2 mutations with clinical characteristics of cancers did not reveal an association of mutations with cancer stages, but p.S464Ter showed a trend toward decreased OS of patients with the mutations (Figure 4D) in pan-cancer and in the individual cancers with >1 p.S464Ter occurrence in informative samples.

**Figure 4.**
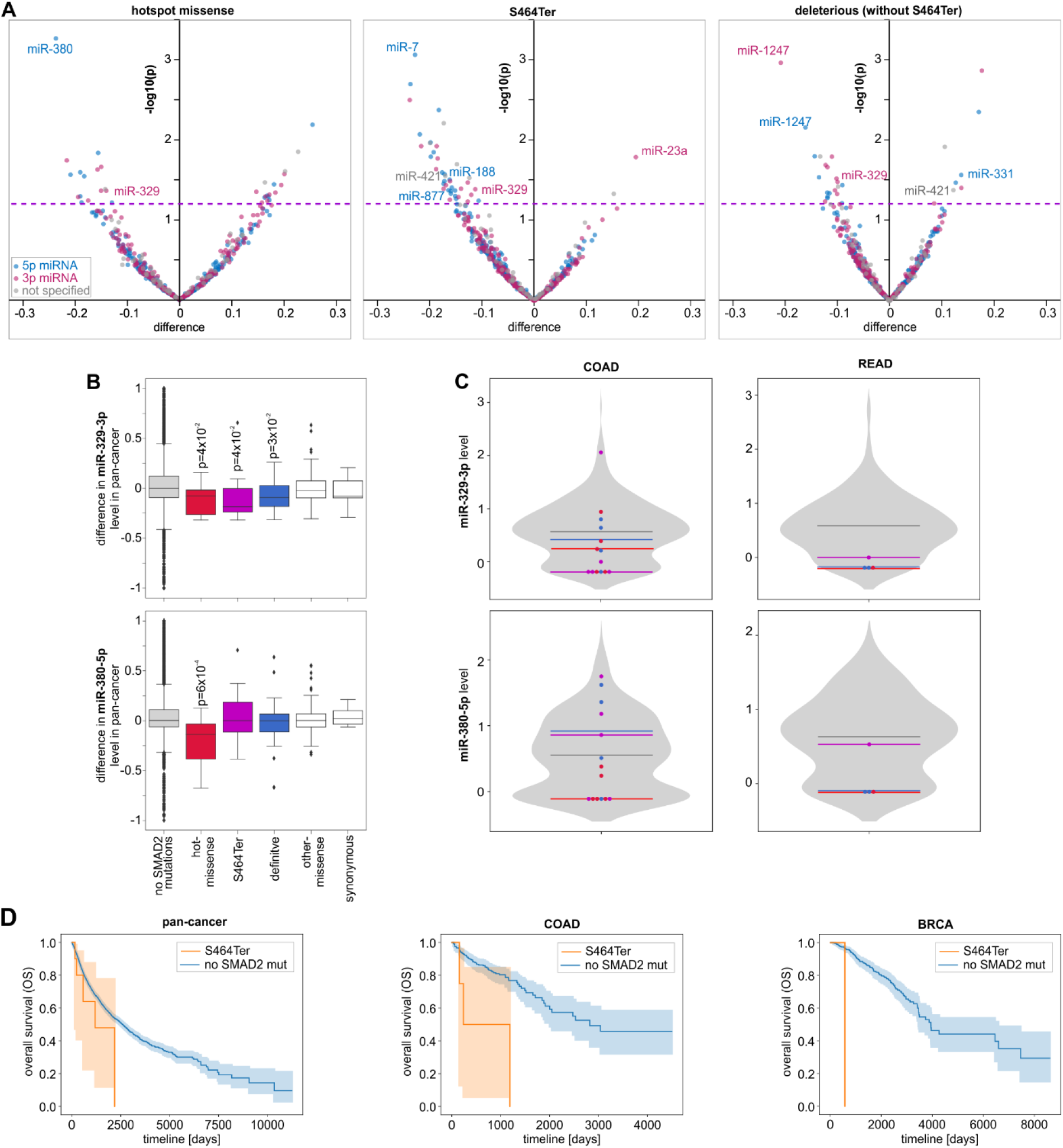
Characteristics of SMAD2 mutations. A) Volcano plots depicting miRNA level alterations in samples with the particular types of *SMAD2* mutations (indicated above the graphs) compared to samples without any *SMAD2* mutations (graph scheme as in Figure 3D). B) Boxplots showing the distribution of the selected miRNA levels (x-axes) in samples with different types of *SMAD2* mutations vs. samples without *SMAD2* mutations. C) Violin plots showing the distribution of non-pan-cancer-normalized levels (y-axis) of the miRNAs shown in B in samples without *SMAD2* mutations and specific samples with different types of *SMAD2* mutations (graph scheme as in Figure 3F). Kaplan-Meier plots showing the OS of patients with p.S464Ter and without any SMAD2 mutations in pan-cancer, COAD, and BRCA.

### Functional consequences of *DICER1* mutations

Another highly mutated gene with several characteristic hotspots is *DICER1*, with a total of 272 mutations (46 mut/kbp), including 38 deleterious mutations and 27 missense mutations located in the aforementioned hotspots in the RIIIa (N=6) and RIIIb (N=21) domains. As shown in Figure 5A, the occurrence of DICER1 mutations does not correlate with gene copy number alterations, and only a small fraction of the *DICER1* mutations coincide with gene deletion or amplification. The miRNA expression analysis at the pan-cancer level showed that the hotspot mutations in both the RIIIa and RIIIb domains were associated with the global downregulation of 5p-miRNAs with simultaneous upregulation of 3p-miRNAs (Figure 5B). This effect of 5p-miRNA downregulation was observed before and was explained by the fact that the hotspot mutations in RIIIb but also in RIIIa [65] affect the activity of the RIIIb domain, preventing cleavage of miRNA precursors at the 5p arm and the release of 5p-miRNAs. The increase in 3p-miRNAs was considered an artifact, i.e., an apparent effect counterbalancing the global decrease in 5-miRNAs, resulting from the standard miRNA level normalization procedure that standardized the amount of each miRNA against the total number of miRNA reads. Surprisingly, we observed a similar effect of decreased 5p-miRNAs and increased 3p-miRNAs in samples with deleterious mutations that do not affect RIIIb but are assumed to lead to complete loss of DICER1 (Figure 5 B and C; for comparison, please see the effect of *SMAD4* mutations in which 5p- and 3p-miRNAs are more or less equally distributed between the decreased and increased miRNAs). The effect of the asymmetrical distribution of altered miRNAs was not observed for other nonhotspot missense mutations outside the RNase domains and synonymous mutations for which, as expected, miRNA level changes were very low (Figure 5C). To avoid potential biases associated with pan-cancer normalization, we performed expression analysis separately for UCEC, which is the cancer type with the most frequent mutations in *DICER1* (~9%), with 2 mutations in RIIIa, 10 mutations in RIIIb, and 9 deleterious mutations. Although of lower statistical power, the UCEC analysis showed a similar effect of the hotspot and deleterious mutations on globally decreased levels of 5p-miRNAs and increased levels of 3p-miRNAs (Figure 5B). To directly check whether the increase in 3p-miRNAs in samples with the *DICER1* mutations is an effect of normalization against the total number of miRNA-specific reads, we normalized the miRNA levels against the level of miR-451a used as a reference gene. miR-451a is a miRNA whose biogenesis is not dependent on DICER1 processing [109]; therefore, its level should not be affected by DICER1 mutations. As shown in Figure 5C, normalization against miR-451a in samples with hotspot mutations abolished the effects of neither 5p-miRNA decreases nor 3p-miRNA increases. This indicates that hotspot mutations, as reported previously [60–62,65,92], decrease 5p-miRNA levels, but contrary to previous reports, they also increase the levels of numerous 3p-miRNAs. KEGG pathway enrichment analysis performed with miRPath v3.0 (Supplementary Table S6) showed that upregulated 3p-miRNAs, among others, are strongly associated with various cancer-related processes, such as ‘TGF-beta signaling pathway’, ‘Pathways in cancer’, ‘ErbB signaling pathway’ and ‘Hippo signaling pathway’, which were found among the top ten most significant associations (adjusted p<0.00001). Consistent with the predicted loss-of-function effect of the deleterious mutations, after normalization against miR-451a, the deleterious mutations are associated almost exclusively with a decrease in miRNA levels (Figure 5C and Supplementary Figure S5). Although an excess of 5p-miRNAs is observed among the decreased miRNAs, there is also a substantial fraction (n=52, 26%) of 3p-miRNAs.

**Figure 5.**
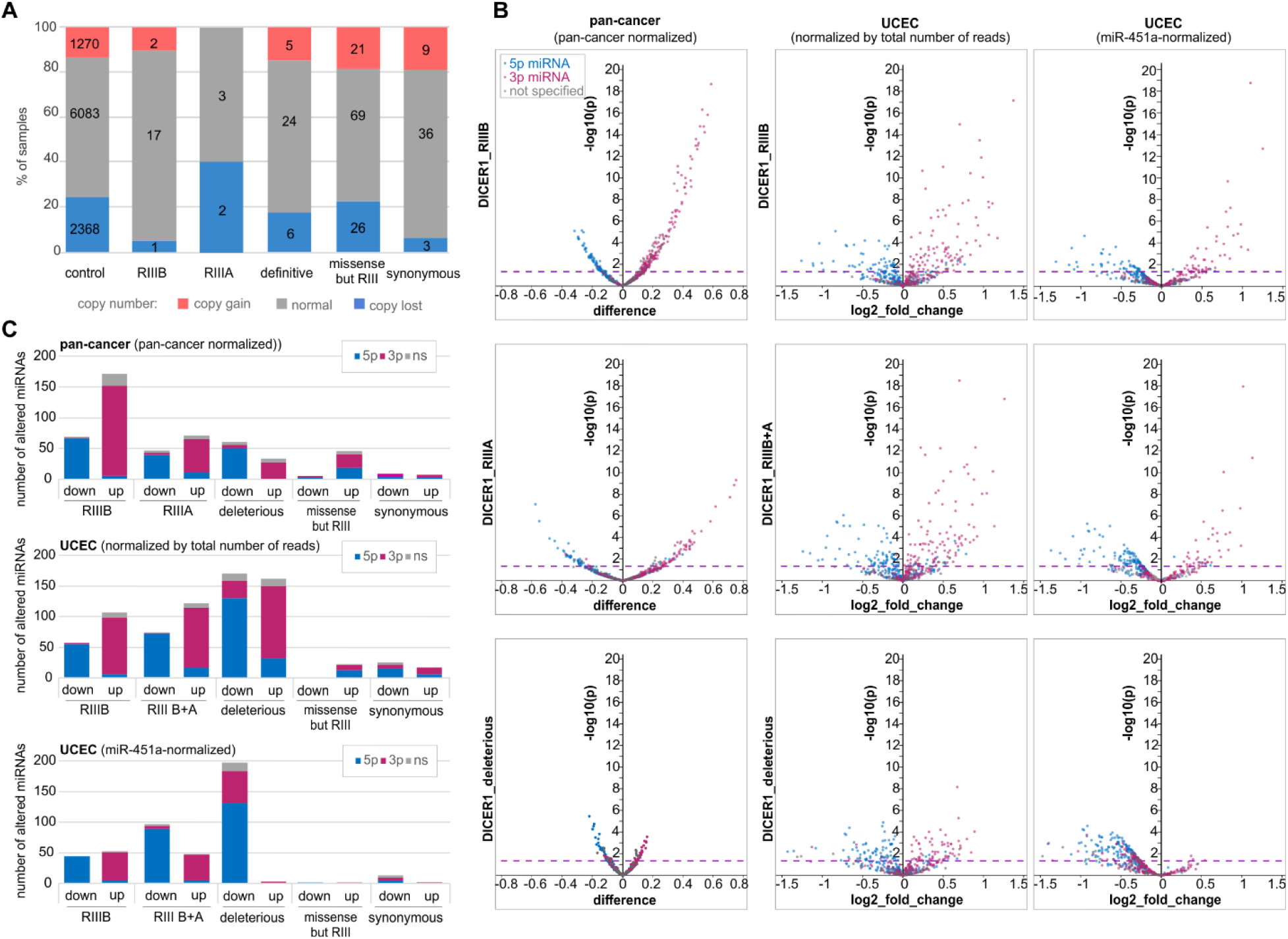
Functional consequences of *DICER1* mutations. A) The proportion of copy number alterations of *DICER1* (y-axis) in samples with different types of *DICER1* mutations. B) Volcano plots depicting miRNA level alterations in pan-cancer (1st column) and UCEC before and after normalization against the miR-451a level (2nd and 3rd columns, respectively) in samples with different types of *DICER1* mutations (indicated on the left). Blue, pink, and gray dots indicate 5p-miRNAs, 3p-miRNAs, and miRNAs whose arm was not specified, respectively. Other graph details as in Figure 3D. C) Each graph shows the proportions of 3p- and 5p-miRNAs (y-axis) among the miRNAs downregulated and upregulated in samples with specified types of *DICER1* mutations (x-axis; at p<0.05). The graphs (from the top) show the alterations in pan-cancer, and UCEC before and after normalization against the miR-451a level.

The lists of miRNAs differentiated in pan-cancer and UCEC with different types of *DICER1* mutations are shown in Supplementary Table S5. Among the altered miRNAs are many miRNAs whose function is well recognized in cancer and many upregulated 3p-miRNAs are passengers of such miRNAs, examples of which are listed in Supplementary Table S7. A comparison of the *DICER1* mutations with clinical characteristics of cancers did not reveal any significant associations.

## DISCUSSION

In this study, we provide a comprehensive pan-cancer analysis of somatic mutagenesis in a panel of 29 miRNA biogenesis genes in 33 adult-onset cancers. In total, we identified 3,649 mutations, composed of 60% missense and 17% deleterious mutations. Overall, approximately 21% of the samples had at least one mutation, but the percentage differed substantially between the cancer types, reaching more than 40% in SKCM or COAD. The most frequently mutated genes were *TNRC6A, DICER1, SMAD4*, and *ZCCHC11*, with mutations in 2-3% of pan-cancer samples. The frequency of mutations in the miRNA biogenesis genes is similar to that in cancer-specific drivers, such as *MYC, ALK, CACNA1A, POLE, BCL2, NOTCH2, MET, HRAS, FGFR*, and *PIK3R1*, with pan-cancer mutation frequencies up to ~3% [110]. Consistent with the potential cancer-specific role of the genes, some of them are much more frequently mutated in particular cancers, for example, *SMAD4* in PAAD (22%), READ (16%), COAD (14%), and ESCA (8%); *DICER1* in SKCM and UCEC (~9%); *TNRC6A* in STAD (9%) and UCEC (7%); ZCCHC6 in ACC (6.5%); *SMAD2* in COAD and READ (~5%); and *PRKRA* in OV (4.3%). The frequencies of mutations in some of the genes are specifically and significantly enriched in particular cancer types.

The high frequency of mutations in *SMAD4* is consistent with the findings of many previous studies in which *SMAD4* was analyzed as a transcription factor playing a role in the activation of many cancer-related genes in response to TGFB/BMP signaling [111–114]. Comprehensive analysis of a large number of samples allowed us to define a profile of *SMAD4* mutations in most common human cancers covered by TCGA. It confirmed the high frequency of *SMAD4* mutations in cancers such as PAAD, COAD, and READ but also revealed a relatively high frequency of the mutations in STAD, LUAD, and ESCA and lower frequencies in many other cancers (e.g., CESC, LUSC, HNC, and CHOL) that are much less studied in the context of *SMAD4* deficiency. With the large collection of mutations, we have identified 8 hotspot residues in the MH2 domain, including some not reported before, and revealed that the proportions of hotspot and deleterious *SMAD4* mutations differ substantially between cancers, with the fraction of hotspot mutations significantly enriched in READ (86%) and the lowest in PAAD (27%). This may suggest different mechanisms and effects of SMAD4 inactivation in different cancer types. We showed that most of the hotspots localize in residues of charged AAs, which may affect the electrostatic interactions of the MH2 domain with corresponding domains in R-SMADs (e.g., SMAD2) critical for SMAD complex formation [85]. Such an effect of both loss-of-charge and gain-of-charge *SMAD4* mutations was computationally predicted in a previous study [115]. Consistently, we showed that all the *SMAD4* hotspots as well as much less frequently mutated hotspot residues in the corresponding MH2 domain in SMAD2 localize at surfaces of the SMAD4:SMAD2 interaction, confirming the role of hotspot mutations in destabilizing the SMAD complex and hence revealing an inhibitory role of these mutations in downstream TGFB/BMP signaling. We showed that both the deleterious and hotspot *SMAD4* mutations frequently (~80%) coincide with gene deletions, which indicates inactivation of the second allele and is consistent with previous observations of frequent LOH in the region [116–119]. Although it was shown that SMAD4 plays a role in the activation of miRNA gene transcription (summarized in [120]), the effect of *SMAD4* mutations on miRNA expression has never been tested. We identified numerous miRNAs differentially expressed in samples with *SMAD4* mutations and showed different patterns of miRNA level changes induced by hotspot and deleterious mutations. While hotspot mutations predominantly downregulate miRNAs, deleterious mutations both increase and decrease miRNA levels. This finding is consistent with the results obtained for *SMAD4* knockdown by RNAi in hepatic stellate cells [73] and again shows a different effect of deleterious mutations that, as a result of NMD, most likely leads to complete loss of SMAD4 and hotspot mutations that modify SMAD4 structure, affecting its interactions with R-SMADs and other proteins. Therefore, the effect of hotspot mutations may result not only from impeded R-SMADs:SMAD4 complex formation but also from a shifted balance in the binding of competing coactivators and corepressors (such as p300/CBP and Ski or SnoN, respectively), whose contribution determines the outcome of signaling events (reviewed in [113,121]).

Similar effects of predominant downregulation of miRNA levels were observed for p.S464Ter, the most frequent *SMAD2* hotspot mutation, as well as for deleterious *SMAD2* mutations. The p.S464Ter mutation localizes in the last exon of *SMAD2* and truncates the last 5 AAs of the protein but most likely does not activate NMD. As the truncated fragment is important for efficient SMAD2 phosphorylation and includes two serine residues (S465 and S467) whose phosphorylation upon TGFB signaling is critical for SMAD4(SMAD2)2 complex formation [91,122,123], the mutation precludes the SMAD complex formation and TGFB/BMP signaling [111]. We showed that the mutation is associated with decreased survival of cancer patients. Of note, two BRCA patients with the mutations showed a strikingly short OS. Regardless of its role in complexes with SMAD4, SMAD2 (along with other R-SMADs) may directly interact with a specific set of miRNA precursors, accelerating their processing by DROSHA [72,124]. This posttranscriptional regulation of miRNA processing may also be affected by SMAD2 mutations, as the MH2 domain of R-SMADs was shown to interact with the P68 RNA helicase participating in the recruitment of DROSHA and DGCR8 to pri-miRNAs [125].

Another gene with characteristic hotspot mutations is *DICER1*. We found 21 hotspot mutations affecting 4 metal-ion-binding AA residues in the RIIIb domain (E1705, D1709, D1810, and E1813) and 6 hotspot mutations in one AA residue (S1344) in RIIIa. These hotspots were previously detected and functionally characterized in various pediatric cancers, including cancers associated with DICER1 syndrome (e.g., pleuropulmonary blastoma) and Wilms’ tumor, as summarized in [54,64]. More recently, they were also investigated in thyroid adenomas [92] and the TCGA cohort, mostly in UCEC samples [65]. It was shown that RIIIb hotspot mutations affect the RIIIb cleavage of the 5p-arm of pre-miRNAs, resulting in inefficient production of 5p-miRNAs [60–62,92]. In a very recent study, it was also shown that the hotspot mutations of the S1344 residue, although located in RIIIa, spatially interfere with RIIIb, resulting in the same effect as that observed for the mutations in RIIIb [65]. The miRNA profiling performed in our study showed a global decrease in 5p-miRNAs and a global increase in 3p-miRNAs in samples with both hotspot and deleterious *DICER1* mutations. A similar effect has been observed before [61,62,65,92,126,127]. The global decrease in 5p-miRNAs has been interpreted as the result of inefficient 5-miRNA processing, but the increase in 3p-miRNAs was considered an artifact of miRNA level normalization (against the total number of miRNA reads) that, as a reflection of the global 5p-miRNA deficit, resulted in a relative increase in unaffected 3p-miRNAs [65]. To eliminate this potential bias, we normalized the miRNA levels against the level of miR-451a, which is a DICER1-independent miRNA whose level is not affected by DICER1 deficiency [109]. However, normalization against the miR-451a level did not eliminate the asymmetrical effect of the hotspot mutations, showing that both the 5p-miRNA decreases and 3p-miRNA increases are real. We have additionally shown that the increased 3p-miRNAs strongly associate with cancer-related terms/pathways and include many miRNAs well recognized in cancer (Supplementary Table S6). The increase in 3p-miRNAs may result from a lack of competition with 5p-miRNAs during transfer to and loading onto the RISC. However, this would require some alternative mechanism of releasing 3p-miRNAs from partially processed (nicked only at 3p-arms) pre-miRNAs and transferring them to the RISC, bypassing the miRNA duplex stage. Although it was previously speculated that AGO2 may play some role in such a process [19,20], in our opinion, this step warrants further investigation. Unlike the hotspot mutations, the deleterious mutations (after normalization against miR-451a) almost exclusively decrease miRNA levels, which confirms their loss-of-function nature. Although the effect of deleterious mutations is more profound for 5p-miRNAs, the mutations also affect 3p-miRNAs (~30%).

With exception of hotspot mutations found in *DICER1*, we found only a few mutations in other miRNA biogenesis genes, i.e., *DROSHA, DGCR8*, and *XPO5*, which have been recently observed in different childhood cancers associated with DICER1 syndrome and in Wilms’ tumor [56–59,66]. This may indicate specific functions of the mutations/genes characteristic of childhood cancers but not playing a role in adult-onset cancers. We also excluded the frequent occurrence of specific indel hotspots in *TRBP* and *XPO5* previously reported at high frequencies in cancers associated with MSI [67,68].

In summary, in this study, we present a comprehensive pan-cancer analysis of somatic mutations accumulated in genes involved in miRNA biogenesis and function. We showed that some of these genes are specifically mutated in particular cancers. We identified many hotspot mutations, including some recurring in specific cancer types. We also extended knowledge about the types and distribution of *SMAD4* mutations and showed their effect on the expression of miRNA genes. We also showed that all hotspot mutations in *SMAD4* and *SMAD2* affect AA residues located at the surface of the SMAD4:SMAD2 interaction. Moreover, we distinguished and further characterized the effects of deleterious and hotspot missense mutations in *DICER1*, among others, showing that hotspot mutations in the RIIIa and RIIIb domains not only decrease the levels of 5p-miRNAs but also increase the levels of 3p-miRNAs. As a result, we have identified numerous miRNAs that are significantly increased or decreased in samples with particular mutation types, including many well-known cancer-related miRNAs. We also linked some of the mutations with the patients’ clinical outcomes. Furthermore, we created a compendium of information presented as an atlas and maps of mutations in miRNA biogenesis genes that may be useful resources of information for studying a particular gene or cancer type.

## ACKNOWLEDGMENTS

The results published here are based upon data generated by the TCGA Research Network: https://www.cancer.gov/tcga (project ID: 16565). This work was supported by research grants from the Polish National Science Centre 2016/22/A/NZ2/00184 (to P.K.).

## AUTHOR CONTRIBUTIONS

PG-M - participated in conceiving the study, participated in writing the scripts, performed most of the computational and statistical analyses, drafted the manuscript, prepared figures, tables, and supplementary materials; MOU-T - participated in conceiving the study, participated in writing the scripts, discussed the study on all steps of analyses, participated in manuscript preparation; PMN - discussed the analyses on all steps of the study, participated in the creation of miRNA biogenesis genes panel; PK - received financing, participated in conceiving the study, supervised and coordinated the study, drafted the manuscript (with PG-M).

---

**Supplementary Figure S1.**
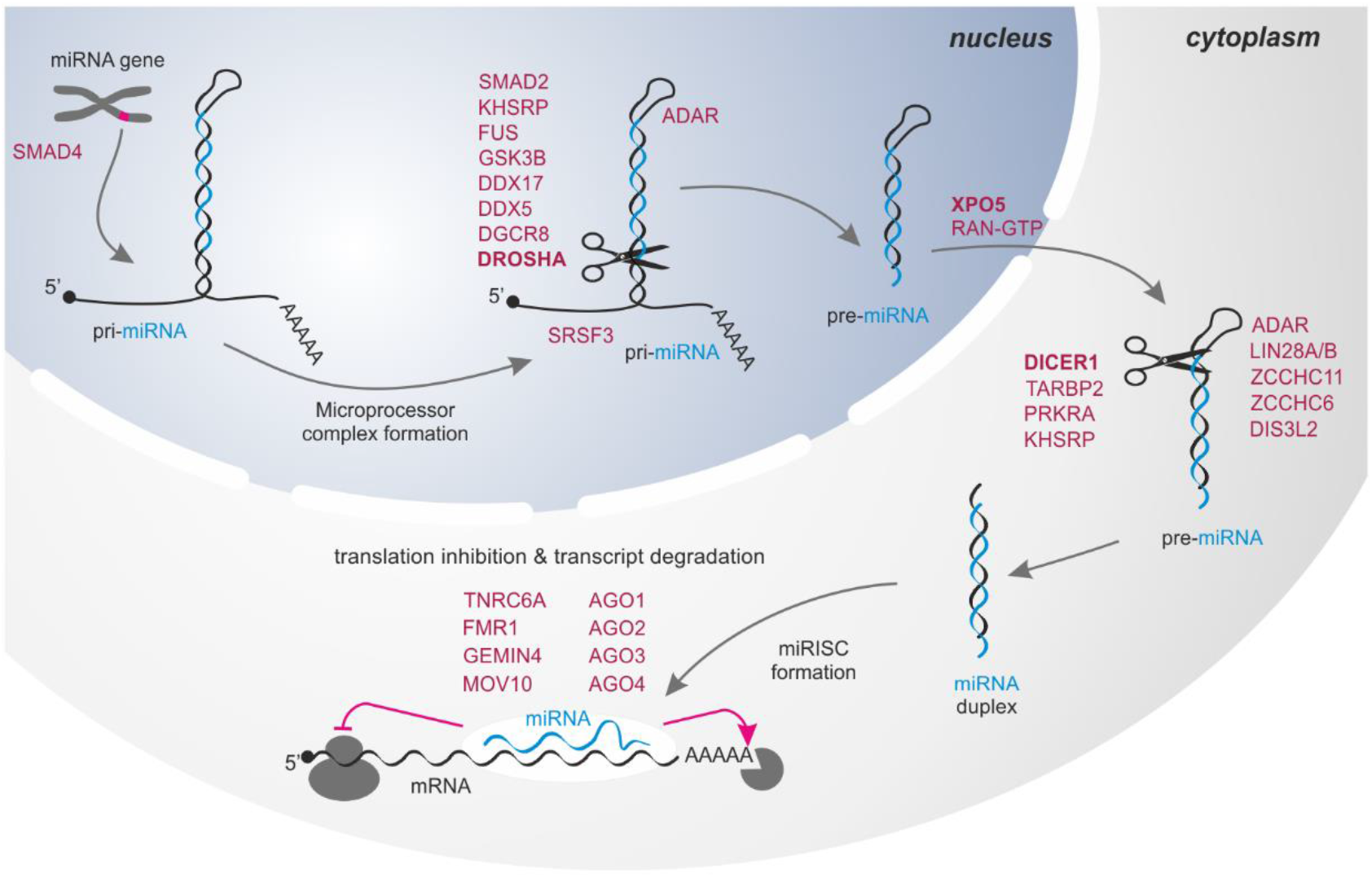
Schematic depiction of miRNA biogenesis and functions of the miRNA biogenesis proteins/genes involved in the subsequent steps.

**Supplementary Figure S2.**
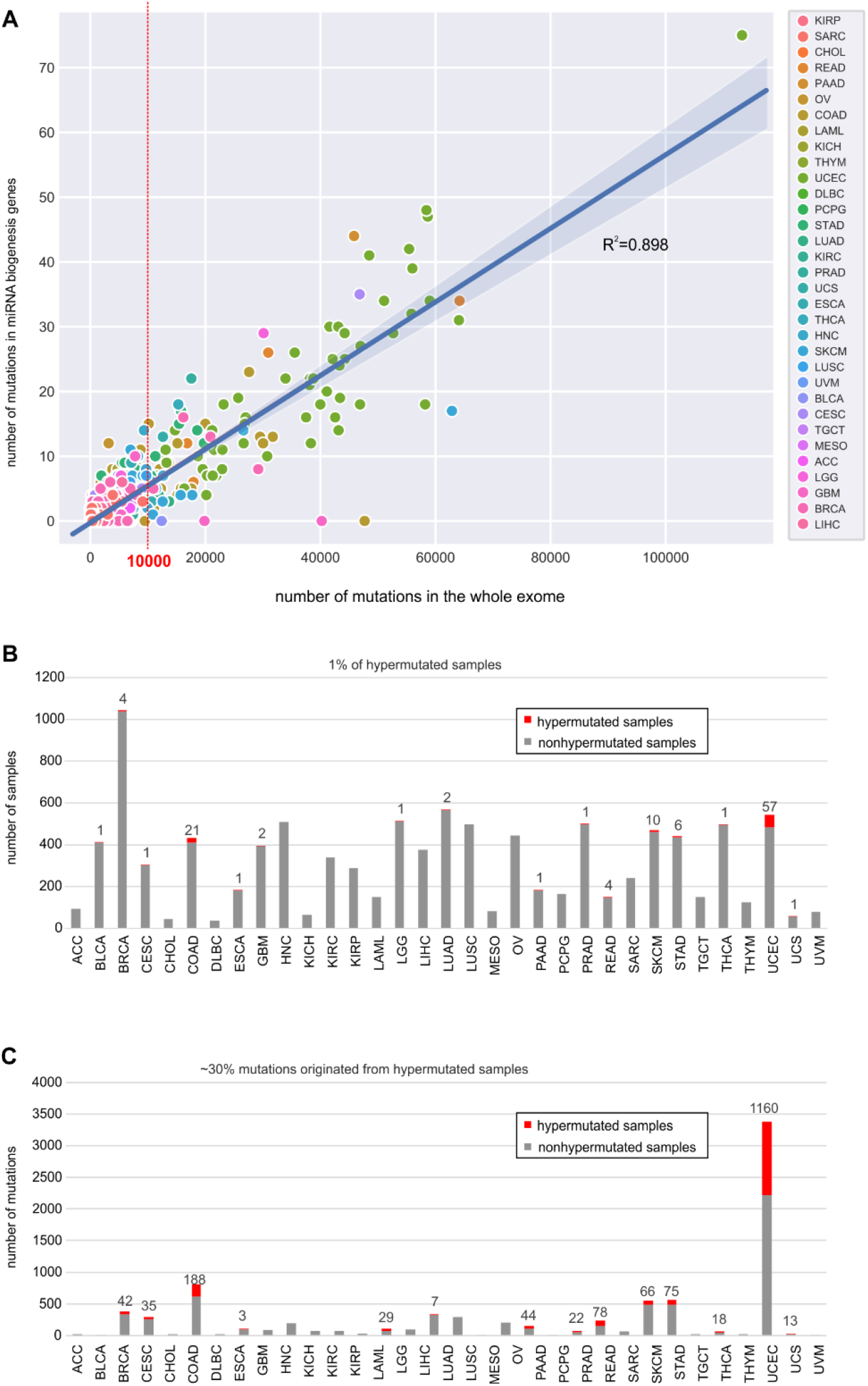
Mutations in hypermutated samples. A) Correlation of the number of mutations in the panel of miRNA biogenesis genes (y-axis) and the general burden of mutations (number of mutations in the whole exome; x-axis) in individual cancer samples. The red vertical line indicates the threshold for hypermutated samples. Cancer types are indicated by different colors. B) The proportions of hypermutated (red bars) and nonhypermutated (gray bars) samples (y-axis) in each cancer type (x-axis). C) The proportions of mutations in hypermutated (red bars) and nonhypermutated (gray bars) samples (y-axis) in each cancer type (x-axis).

**Supplementary Figure S3.**
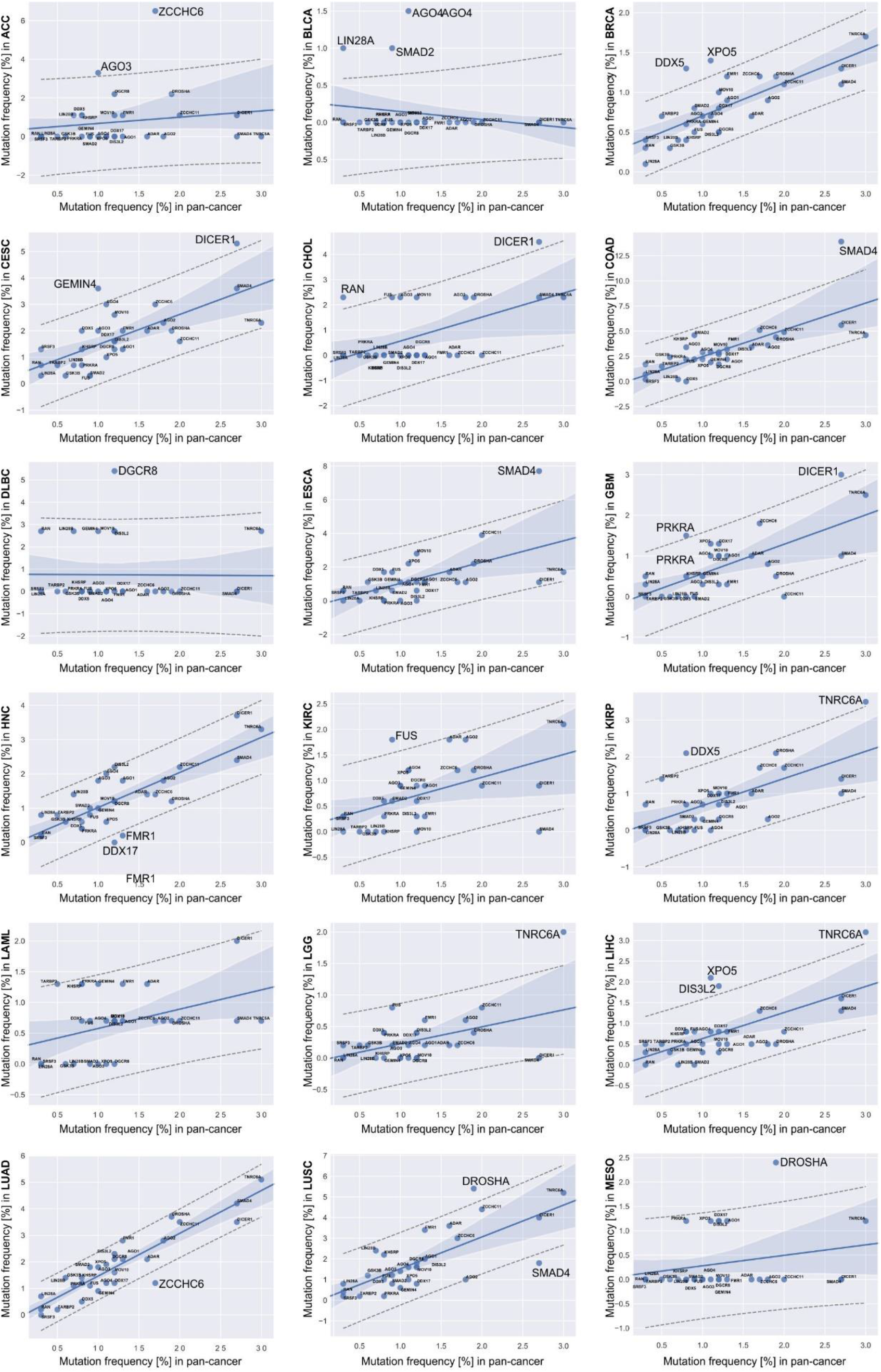
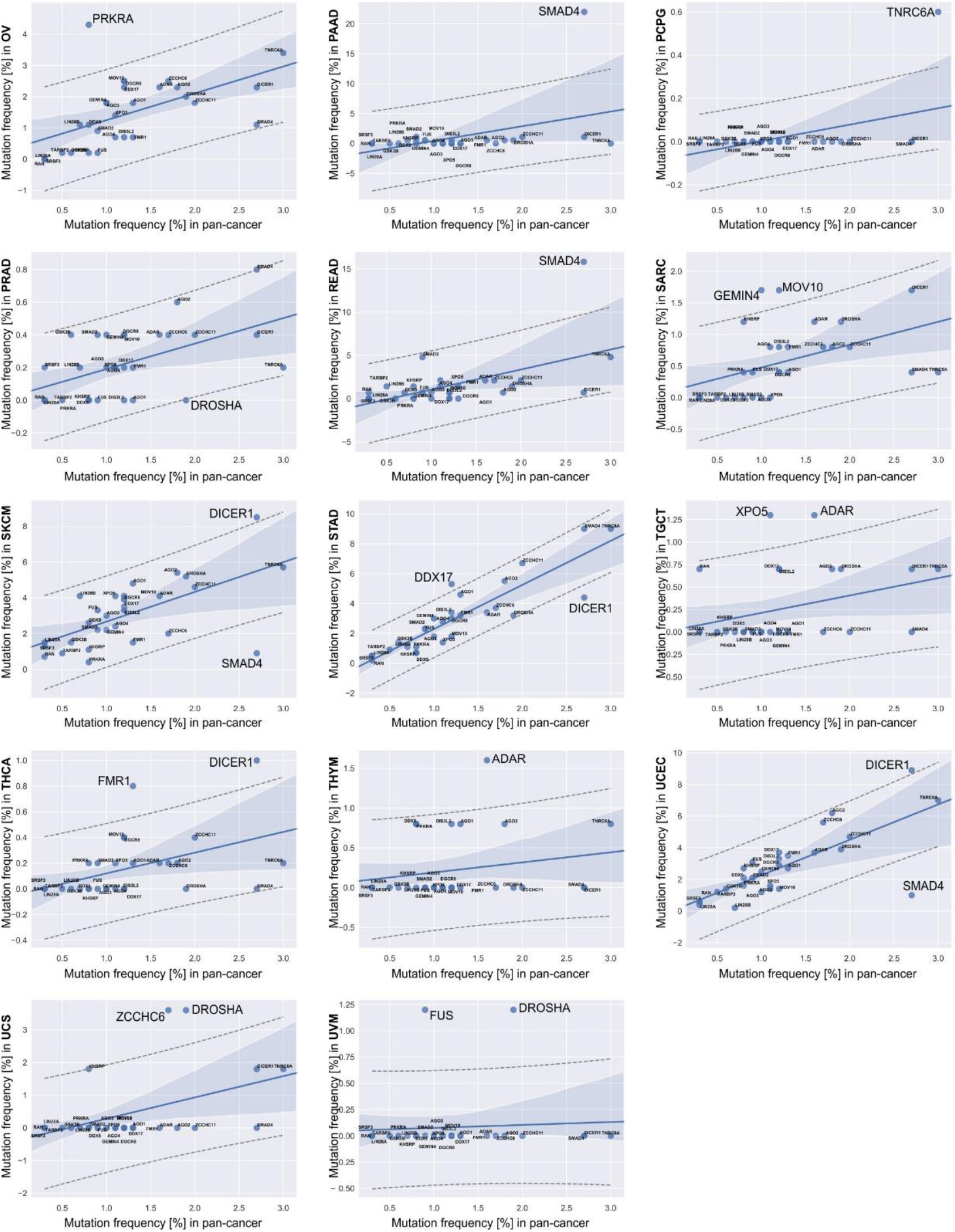
Scatterplots showing the correlation between the frequencies of mutations in the miRNA biogenesis genes observed in the individual cancer types (y-axis) and in the remaining pan-cancer (x-axis) samples. In each graph, dots indicating the particular miRNA biogenesis genes, the regression line, and dashed lines representing the confidence interval with 95% probability are shown. Note that the informativeness of some of the graphs is low due to the very low number of mutations identified in some cancer types.

**Supplementary Figure S4.**
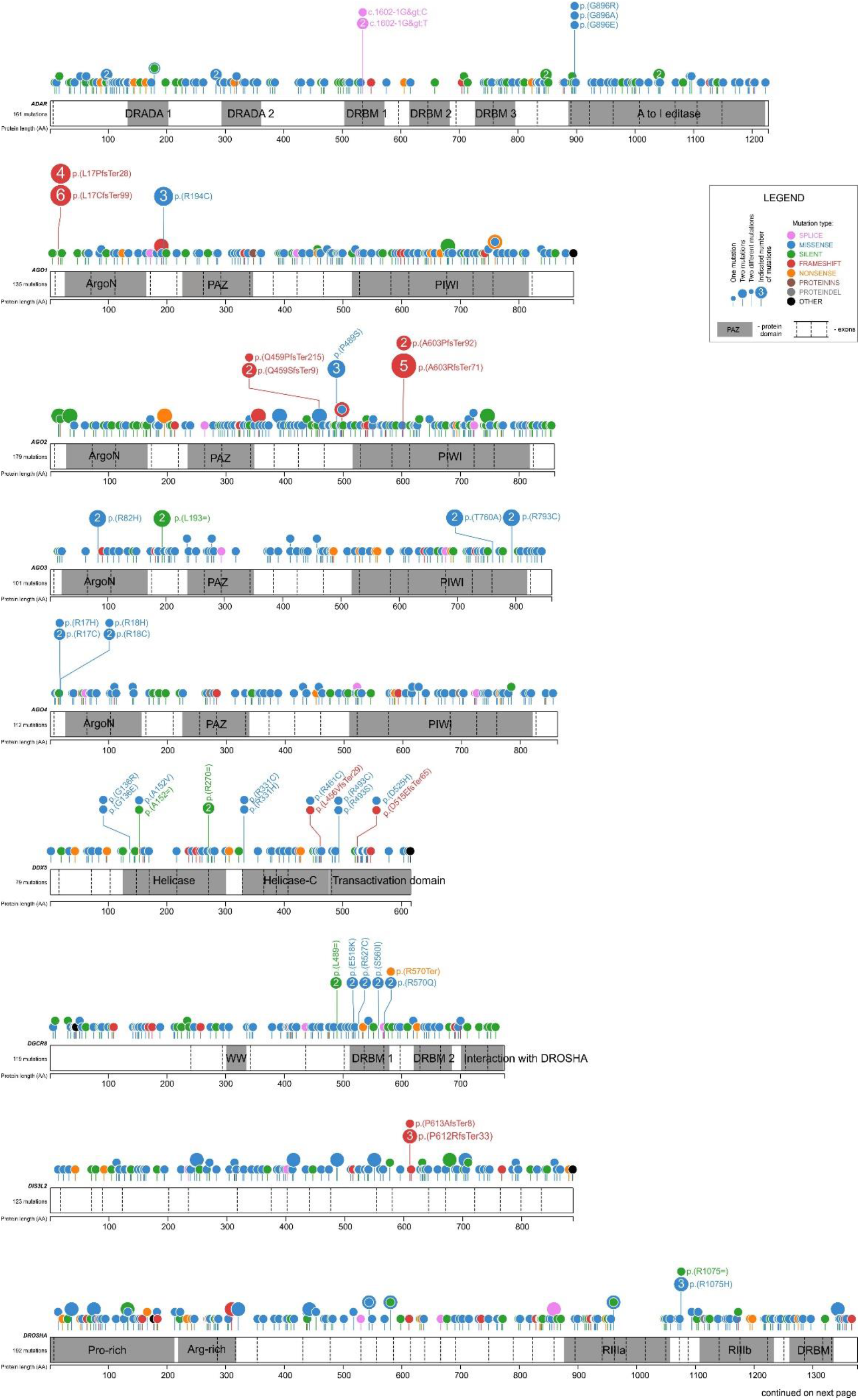
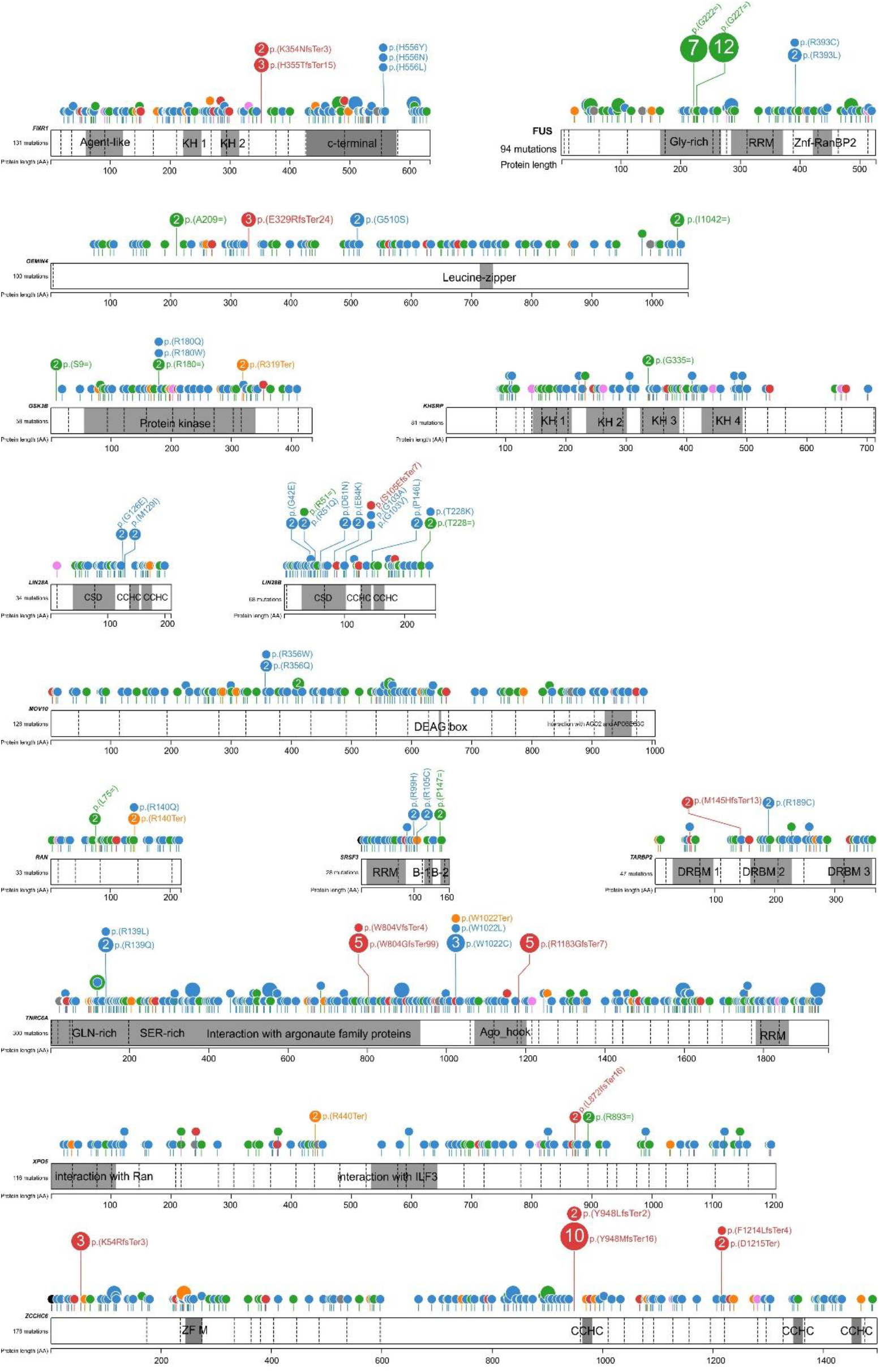
Distribution of the identified mutations in the miRNA biogenesis genes. The scheme of the figure is as shown in Figure 2.

## SUPPLEMENTARY TABLES

**Supplementary Table S1**: Genomic coordinates of 29 miRNA biogenesis genes (hg38) [xlsx]

**Supplementary Table S2:** Summary of TCGA samples and mutations in the panel of miRNA biogenesis genes per cancer type [xlsx]

**Supplementary Table S3:** The mutations identified in the panel of miRNA biogenesis genes in the pan-cancer [xlsx]

**Supplementary Table S4:** Overmutation of the particular miRNA biogenesis genes in specific cancer types (p-values) [xlsx]

**Supplementary Table S5:** Lists of altered miRNAs in samples with the specified type of mutations in *SMAD4*, *SMAD2*, and *DICER1* [xlsx]

**Supplementary Table S6:** List of pathways associated with panels of miRNAs either decreased or increased by the specified type of mutations in *SMAD4* and *DICER1* [xlsx]

**Supplementary Table S7** The list of miRNAs well recognized in cancer that were differentiated by different types of *DICER1* mutations [xlsx]

## REFERENCES

1. Calin GA, Dumitru CD, Shimizu M, Bichi R, Zupo S, Noch E, et al. Frequent deletions and down-regulation of micro-RNA genes miR15 and miR16 at 13q14 in chronic lymphocytic leukemia. Proc Natl Acad Sci U S A. 2002;99:15524–9.

2. Peng Y, Croce CM. The role of MicroRNAs in human cancer. Signal Transduct Target Ther. 2016;1:15004.

3. Lu J, Getz G, Miska EA, Alvarez-Saavedra E, Lamb J, Peck D, et al. MicroRNA expression profiles classify human cancers. Nature. 2005;435:834–8.

4. Volinia S, Calin GA, Liu C-G, Ambs S, Cimmino A, Petrocca F, et al. A microRNA expression signature of human solid tumors defines cancer gene targets. Proc Natl Acad Sci U S A. 2006;103:2257–61.

5. Esquela-Kerscher A, Slack FJ. Oncomirs -microRNAs with a role in cancer. Nat Rev Cancer. 2006;6:259–69.

6. Tan W, Liu B, Qu S, Liang G, Luo W, Gong C. MicroRNAs and cancer: Key paradigms in molecular therapy. Oncol Lett. 2018;15:2735–42.

7. Krutovskikh VA, Herceg Z. Oncogenic microRNAs (OncomiRs) as a new class of cancer biomarkers. BioEssays News Rev Mol Cell Dev Biol. 2010;32:894–904.

8. Kasinski AL, Slack FJ. Epigenetics and genetics. MicroRNAs en route to the clinic: progress in validating and targeting microRNAs for cancer therapy. Nat Rev Cancer. 2011; 11:849–64.

9. Cui M, Wang H, Yao X, Zhang D, Xie Y, Cui R, et al. Circulating MicroRNAs in Cancer: Potential and Challenge. Front Genet. 2019;10:626.

10. Bartel DP. MicroRNAs: genomics, biogenesis, mechanism, and function. Cell. 2004;116:281–97.

11. Ha M, Kim VN. Regulation of microRNA biogenesis. Nat Rev Mol Cell Biol. 2014;15:509–24.

12. Kim VN, Han J, Siomi MC. Biogenesis of small RNAs in animals. Nat Rev Mol Cell Biol. 2009;10:126–39.

13. Bartel DP. Metazoan MicroRNAs. Cell. 2018;173:20–51.

14. Michlewski G, Cáceres JF. Post-transcriptional control of miRNA biogenesis. RNA N Y N. 2019;25:1–16.

15. Treiber T, Treiber N, Meister G. Regulation of microRNA biogenesis and its crosstalk with other cellular pathways. Nat Rev Mol Cell Biol. 2019;20:5–20.

16. Ustianenko D, Hrossova D, Potesil D, Chalupnikova K, Hrazdilova K, Pachernik J, et al. Mammalian DIS3L2 exoribonuclease targets the uridylated precursors of let-7 miRNAs. RNA N Y N. 2013;19:1632–8.

17. Heo I, Joo C, Kim Y-K, Ha M, Yoon M-J, Cho J, et al. TUT4 in concert with Lin28 suppresses microRNA biogenesis through pre-microRNA uridylation. Cell. 2009;138:696–708.

18. Sibley CR, Seow Y, Saayman S, Dijkstra KK, El Andaloussi S, Weinberg MS, et al. The biogenesis and characterization of mammalian microRNAs of mirtron origin. Nucleic Acids Res. 2012;40:438–48.

19. Cheloufi S, Dos Santos CO, Chong MMW, Hannon GJ. A dicer-independent miRNA biogenesis pathway that requires Ago catalysis. Nature. 2010;465:584–9.

20. Cifuentes D, Xue H, Taylor DW, Patnode H, Mishima Y, Cheloufi S, et al. A novel miRNA processing pathway independent of Dicer requires Argonaute2 catalytic activity. Science. 2010;328:1694–8.

21. Kozomara A, Griffiths-Jones S. miRBase: annotating high confidence microRNAs using deep sequencing data. Nucleic Acids Res. 2014;42:D68–73.

22. Kozomara A, Birgaoanu M, Griffiths-Jones S. miRBase: from microRNA sequences to function. Nucleic Acids Res. 2019;47:D155–62.

23. Chou C-H, Shrestha S, Yang C-D, Chang N-W, Lin Y-L, Liao K-W, et al. miRTarBase update 2018: a resource for experimentally validated microRNA-target interactions. Nucleic Acids Res. 2018;46:D296–302.

24. Friedman RC, Farh KK-H, Burge CB, Bartel DP. Most mammalian mRNAs are conserved targets of microRNAs. Genome Res. 2009;19:92–105.

25. Kong W, Yang H, He L, Zhao J, Coppola D, Dalton WS, et al. MicroRNA-155 is regulated by the transforming growth factor beta/Smad pathway and contributes to epithelial cell plasticity by targeting RhoA. Mol Cell Biol. 2008;28:6773–84.

26. Tong Z, Cui Q, Wang J, Zhou Y. TransmiR v2.0: an updated transcription factor-microRNA regulation database. Nucleic Acids Res. 2019;47:D253–8.

27. Kato M, Putta S, Wang M, Yuan H, Lanting L, Nair I, et al. TGF-beta activates Akt kinase through a microRNA-dependent amplifying circuit targeting PTEN. Nat Cell Biol. 2009;11:881–9.

28. Chung ACK, Huang XR, Meng X, Lan HY. miR-192 mediates TGF-beta/Smad3-driven renal fibrosis. J Am Soc Nephrol JASN. 2010;21:1317–25.

29. Czubak K, Lewandowska MA, Klonowska K, Roszkowski K, Kowalewski J, Figlerowicz M, et al. High copy number variation of cancer-related microRNA genes and frequent amplification of DICER1 and DROSHA in lung cancer. Oncotarget. 2015;6:23399–416.

30. Calin GA, Sevignani C, Dumitru CD, Hyslop T, Noch E, Yendamuri S, et al. Human microRNA genes are frequently located at fragile sites and genomic regions involved in cancers. Proc Natl Acad Sci U S A. 2004;101:2999–3004.

31. Zhang L, Huang J, Yang N, Greshock J, Megraw MS, Giannakakis A, et al. microRNAs exhibit high frequency genomic alterations in human cancer. Proc Natl Acad Sci U S A. 2006;103:9136–41.

32. Fazi F, Racanicchi S, Zardo G, Starnes LM, Mancini M, Travaglini L, et al. Epigenetic silencing of the myelopoiesis regulator microRNA-223 by the AML1/ETO oncoprotein. Cancer Cell. 2007;12:457–66.

33. Han L, Witmer PD, Casey E, Valle D, Sukumar S. DNA methylation regulates MicroRNA expression. Cancer Biol Ther. 2007;6:1284–8.

34. Saito Y, Jones PA. Epigenetic activation of tumor suppressor microRNAs in human cancer cells. Cell Cycle Georget Tex. 2006;5:2220–2.

35. Ali Syeda Z, Langden SSS, Munkhzul C, Lee M, Song SJ. Regulatory Mechanism of MicroRNA Expression in Cancer. Int J Mol Sci. 2020;21.

36. Lujambio A, Calin GA, Villanueva A, Ropero S, Sánchez-Céspedes M, Blanco D, et al. A microRNA DNA methylation signature for human cancer metastasis. Proc Natl Acad Sci U S A. 2008;105:13556–61.

37. Lujambio A, Esteller M. How epigenetics can explain human metastasis: a new role for microRNAs. Cell Cycle Georget Tex. 2009;8:377–82.

38. Glaich O, Parikh S, Bell RE, Mekahel K, Donyo M, Leader Y, et al. DNA methylation directs microRNA biogenesis in mammalian cells. Nat Commun. Nature Publishing Group; 2019;10:1–11.

39. Alarcón CR, Lee H, Goodarzi H, Halberg N, Tavazoie SF. N6-methyl-adenosine (m6A) marks primary microRNAs for processing. Nature. 2015;519:482–5.

40. Konno M, Koseki J, Asai A, Yamagata A, Shimamura T, Motooka D, et al. Distinct methylation levels of mature microRNAs in gastrointestinal cancers. Nat Commun. 2019;10:3888.

41. Saunders MA, Liang H, Li W-H. Human polymorphism at microRNAs and microRNA target sites. Proc Natl Acad Sci U S A. 2007;104:3300–5.

42. Marcinkowska M, Szymanski M, Krzyzosiak WJ, Kozlowski P. Copy number variation of microRNA genes in the human genome. BMC Genomics. 2011;12:183.

43. Króliczewski J, Sobolewska A, Lejnowski D, Collawn JF, Bartoszewski R. microRNA single polynucleotide polymorphism influences on microRNA biogenesis and mRNA target specificity. Gene. 2018;640:66–72.

44. Sun G, Yan J, Noltner K, Feng J, Li H, Sarkis DA, et al. SNPs in human miRNA genes affect biogenesis and function. RNA N Y N. 2009;15:1640–51.

45. Galka-Marciniak P, Urbanek-Trzeciak MO, Nawrocka PM, Dutkiewicz A, Giefing M, Lewandowska MA, et al. Somatic Mutations in miRNA Genes in Lung Cancer-Potential Functional Consequences of Non-Coding Sequence Variants. Cancers. 2019;11.

46. Oak N, Ghosh R, Huang K-L, Wheeler DA, Ding L, Plon SE. Framework for microRNA variant annotation and prioritization using human population and disease datasets. Hum Mutat. 2019;40:73–89.

47. Bhattacharya A, Cui Y. SomamiR 2.0: a database of cancer somatic mutations altering microRNA-ceRNA interactions. Nucleic Acids Res. 2016;44:D1005–1010.

48. Hornshøj H, Nielsen MM, Sinnott-Armstrong NA, Świtnicki MP, Juul M, Madsen T, et al. Pan-cancer screen for mutations in non-coding elements with conservation and cancer specificity reveals correlations with expression and survival. NPJ Genomic Med. 2018;3:1.

49. Urbanek-Trzeciak MO, Galka-Marciniak P, Nawrocka PM, Kowal E, Szwec S, Giefing M, et al. Pan-Cancer analysis of somatic mutations in miRNA genes. bioRxiv. Cold Spring Harbor Laboratory; 2020;2020.06.05.136036.

50. Tay Y, Rinn J, Pandolfi PP. The multilayered complexity of ceRNA crosstalk and competition. Nature. 2014;505:344–52.

51. Poliseno L, Salmena L, Zhang J, Carver B, Haveman WJ, Pandolfi PP. A coding-independent function of gene and pseudogene mRNAs regulates tumour biology. Nature. 2010;465:1033–8.

52. Joyce BT, Zheng Y, Zhang Z, Liu L, Kocherginsky M, Murphy R, et al. miRNA-Processing Gene Methylation and Cancer Risk. Cancer Epidemiol Biomark Prev Publ Am Assoc Cancer Res Cosponsored Am Soc Prev Oncol. 2018;27:550–7.

53. Merritt WM, Lin YG, Han LY, Kamat AA, Spannuth WA, Schmandt R, et al. Dicer, Drosha, and Outcomes in Patients with Ovarian Cancer. N Engl J Med. 2008;359:2641–50.

54. Foulkes WD, Priest JR, Duchaine TF. DICER1: mutations, microRNAs and mechanisms. Nat Rev Cancer. 2014;14:662–72.

55. Robertson JC, Jorcyk CL, Oxford JT. DICER1 Syndrome: DICER1 Mutations in Rare Cancers. Cancers [Internet]. 2018 [cited 2020 Jun 12];10. Available from: https://www.ncbi.nlm.nih.gov/pmc/articles/PMC5977116/

56. Torrezan GT, Ferreira EN, Nakahata AM, Barros BDF, Castro MTM, Correa BR, et al. Recurrent somatic mutation in DROSHA induces microRNA profile changes in Wilms tumour. Nat Commun. Nature Publishing Group; 2014;5:1–10.

57. Walz AL, Ooms A, Gadd S, Gerhard DS, Smith MA, Guidry Auvil JM, et al. Recurrent DGCR8, DROSHA, and SIX Homeodomain Mutations in Favorable Histology Wilms Tumors. Cancer Cell. 2015;27:286–97.

58. Wegert J, Ishaque N, Vardapour R, Geörg C, Gu Z, Bieg M, et al. Mutations in the SIX1/2 pathway and the DROSHA/DGCR8 miRNA microprocessor complex underlie high-risk blastemal type Wilms tumors. Cancer Cell. 2015;27:298–311.

59. Rakheja D, Chen KS, Liu Y, Shukla AA, Schmid V, Chang T-C, et al. Somatic mutations in DROSHA and DICER1 impair microRNA biogenesis through distinct mechanisms in Wilms tumours. Nat Commun. Nature Publishing Group; 2014;5:1–11.

60. Heravi-Moussavi A, Anglesio MS, Cheng S-WG, Senz J, Yang W, Prentice L, et al. Recurrent somatic DICER1 mutations in nonepithelial ovarian cancers. N Engl J Med. 2012;366:234–42.

61. Anglesio MS, Wang Y, Yang W, Senz J, Wan A, Heravi-Moussavi A, et al. Cancer-associated somatic DICER1 hotspot mutations cause defective miRNA processing and reverse-strand expression bias to predominantly mature 3p strands through loss of 5p strand cleavage. J Pathol. 2013;229:400–9.

62. Gurtan AM, Lu V, Bhutkar A, Sharp PA. In vivo structure-function analysis of human Dicer reveals directional processing of precursor miRNAs. RNA N Y N. 2012;18:1116–22.

63. Kurzynska-Kokorniak A, Koralewska N, Pokornowska M, Urbanowicz A, Tworak A, Mickiewicz A, et al. The many faces of Dicer: the complexity of the mechanisms regulating Dicer gene expression and enzyme activities. Nucleic Acids Res. 2015;43:4365–80.

64. Lin S, Gregory RI. MicroRNA biogenesis pathways in cancer. Nat Rev Cancer. 2015;15:321–33.

65. Vedanayagam J, Chatila WK, Aksoy BA, Majumdar S, Skanderup AJ, Demir E, et al. Cancer-associated mutations in DICER1 RNase IIIa and IIIb domains exert similar effects on miRNA biogenesis. Nat Commun. 2019;10:3682.

66. Gadd S, Huff V, Walz AL, Ooms AHAG, Armstrong AE, Gerhard DS, et al. A Children’s Oncology Group and TARGET initiative exploring the genetic landscape of Wilms tumor. Nat Genet. 2017;49:1487–94.

67. Melo SA, Moutinho C, Ropero S, Calin GA, Rossi S, Spizzo R, et al. A Genetic Defect in Exportin-5 Traps Precursor MicroRNAs in the Nucleus of Cancer Cells. Cancer Cell. 2010;18:303–15.

68. Melo SA, Ropero S, Moutinho C, Aaltonen LA, Yamamoto H, Calin GA, et al. A TARBP2 mutation in human cancer impairs microRNA processing and DICER1 function. Nat Genet. Nature Publishing Group; 2009;41:365–70.

69. Yang G, Yang X. Smad4-mediated TGF-β signaling in tumorigenesis. Int J Biol Sci. 2010;6:1–8.

70. Massagué J, Chen YG. Controlling TGF-beta signaling. Genes Dev. 2000;14:627–44.

71. Zhao M, Mishra L, Deng C-X. The role of TGF-β/SMAD4 signaling in cancer. Int J Biol Sci. 2018;14:111–23.

72. Davis BN, Hilyard AC, Nguyen PH, Lagna G, Hata A. Smad proteins bind a conserved RNA sequence to promote microRNA maturation by Drosha. Mol Cell. 2010;39:373–84.

73. Davoodian P, Ravanshad M, Hosseini SY, Khanizadeh S, Almasian M, Nejati Zadeh A, et al. Effect of TGF-β/smad signaling pathway blocking on expression profiles of miR-335, miR-150, miR-194, miR-27a, and miR-199a of hepatic stellate cells (HSCs). Gastroenterol Hepatol Bed Bench. 2017;10:112–7.

74. Song X, Zhong H, Wu Q, Wang M, Zhou J, Zhou Y, et al. Association between SNPs in microRNA machinery genes and gastric cancer susceptibility, invasion, and metastasis in Chinese Han population. Oncotarget. 2017;8:86435–46.

75. Xie Y, Wang Y, Zhao Y, Guo Z. Single-nucleotide polymorphisms of microRNA processing machinery genes are associated with risk for gastric cancer. OncoTargets Ther. 2015;8:567–71.

76. Li X, Tian X, Zhang B, Zhang Y, Chen J. Variation in dicer gene is associated with increased survival in T-cell lymphoma. PloS One. 2012;7:e51640.

77. Nikolić Z, Savić Pavićević D, Vučić N, Cerović S, Vukotić V, Brajušković G. Genetic variants in RNA-induced silencing complex genes and prostate cancer. World J Urol. 2017;35:613–24.

78. Qian F, Feng Y, Zheng Y, Ogundiran TO, Ojengbede O, Zheng W, et al. Genetic variants in microRNA and microRNA biogenesis pathway genes and breast cancer risk among women of African ancestry. Hum Genet. 2016;135:1145–59.

79. Sung H, Jeon S, Lee K-M, Han S, Song M, Choi J-Y, et al. Common genetic polymorphisms of microRNA biogenesis pathway genes and breast cancer survival. BMC Cancer. 2012;12:195.

80. Yang H, Dinney CP, Ye Y, Zhu Y, Grossman HB, Wu X. Evaluation of genetic variants in microRNA-related genes and risk of bladder cancer. Cancer Res. 2008;68:2530–7.

81. Liang D, Meyer L, Chang DW, Lin J, Pu X, Ye Y, et al. Genetic variants in MicroRNA biosynthesis pathways and binding sites modify ovarian cancer risk, survival, and treatment response. Cancer Res. 2010;70:9765–76.

82. Liu J, Liu J, Wei M, He Y, Liao B, Liao G, et al. Genetic Variants in the MicroRNA Machinery Gene GEMIN4 Are Associated with Risk of Prostate Cancer: A Case-control Study of the Chinese Han Population. DNA Cell Biol. Mary Ann Liebert, Inc.; 2012;31:1296.

83. Mullany LE, Herrick JS, Wolff RK, Buas MF, Slattery ML. Impact of polymorphisms in microRNA biogenesis genes on colon cancer risk and microRNA expression levels: a population-based, case-control study. BMC Med Genomics. 2016;9:21.

84. Fang X, Yin Z, Li X, Xia L, Zhou B. Polymorphisms in GEMIN4 and AGO1 Genes Are Associated with the Risk of Lung Cancer: A Case-Control Study in Chinese Female Non-Smokers. Int J Environ Res Public Health. 2016;13.

85. Chacko BM, Qin BY, Tiwari A, Shi G, Lam S, Hayward LJ, et al. Structural basis of heteromeric smad protein assembly in TGF-beta signaling. Mol Cell. 2004;15:813–23.

86. McLaren W, Gil L, Hunt SE, Riat HS, Ritchie GRS, Thormann A, et al. The Ensembl Variant Effect Predictor. Genome Biol. 2016;17:122.

87. Zhou X, Edmonson MN, Wilkinson MR, Patel A, Wu G, Liu Y, et al. Exploring genomic alteration in pediatric cancer using ProteinPaint. Nat Genet. 2016;48:4–6.

88. UniProt Consortium. UniProt: a worldwide hub of protein knowledge. Nucleic Acids Res. 2019;47:D506–15.

89. Cameron Davidson-Pilon, Jonas Kalderstam, Noah Jacobson, Paul Zivich, Ben Kuhn, Mike Williamson, et al. CamDavidsonPilon/lifelines: v0.24.8 [Internet]. Zenodo; 2020 [cited 2020 Jun 2]. Available from: https://zenodo.org/record/3833188

90. Lawrence MS, Stojanov P, Polak P, Kryukov GV, Cibulskis K, Sivachenko A, et al. Mutational heterogeneity in cancer and the search for new cancer-associated genes. Nature. 2013;499:214–8.

91. Abdollah S, Macías-Silva M, Tsukazaki T, Hayashi H, Attisano L, Wrana JL. TbetaRI phosphorylation of Smad2 on Ser465 and Ser467 is required for Smad2-Smad4 complex formation and signaling. J Biol Chem. 1997;272:27678–85.

92. Poma AM, Condello V, Denaro M, Torregrossa L, Elisei R, Vitti P, et al. DICER1 somatic mutations strongly impair miRNA processing even in benign thyroid lesions. Oncotarget. 2019;10:1785–97.

93. Macias MJ, Martin-Malpartida P, Massagué J. Structural determinants of SMAD function in TGF-β signaling. Trends Biochem Sci. 2015;40:296–308.

94. Holtzhausen A, Golzio C, How T, Lee Y-H, Schiemann WP, Katsanis N, et al. Novel bone morphogenetic protein signaling through Smad2 and Smad3 to regulate cancer progression and development. FASEB J. 2014;28:1248–67.

95. Qin W, Chung ACK, Huang XR, Meng X-M, Hui DSC, Yu C-M, et al. TGF-β/Smad3 signaling promotes renal fibrosis by inhibiting miR-29. J Am Soc Nephrol JASN. 2011;22:1462–74.

96. Wu C-W, Storey KB. Regulation of Smad mediated microRNA transcriptional response in ground squirrels during hibernation. Mol Cell Biochem. 2018;439:151–61.

97. Wang N, Tan H-Y, Feng Y-G, Zhang C, Chen F, Feng Y. microRNA-23a in Human Cancer: Its Roles, Mechanisms and Therapeutic Relevance. Cancers [Internet]. 2018 [cited 2020 Apr 23];11. Available from: https://www.ncbi.nlm.nih.gov/pmc/articles/PMC6356664/

98. Rawat VPS, Götze M, Rasalkar A, Vegi NM, Ihme S, Thoene S, et al. The microRNA miR-196b acts as tumor suppressor in Cdx2 driven acute myeloid leukemia. Haematologica. 2019;

99. Liang G, Meng W, Huang X, Zhu W, Yin C, Wang C, et al. miR-196b-5p-mediated downregulation of TSPAN12 and GATA6 promotes tumor progression in non-small cell lung cancer. Proc Natl Acad Sci U S A. 2020;117:4347–57.

100. Liang J, Zhou W, Sakre N, DeVecchio J, Ferrandon S, Ting AH, et al. Epigenetically regulated miR-1247 functions as a novel tumour suppressor via MYCBP2 in methylator colon cancers. Br J Cancer. 2018;119:1267–77.

101. Zeng B, Li Y, Feng Y, Lu M, Yuan H, Yi Z, et al. Downregulated miR-1247-5p associates with poor prognosis and facilitates tumor cell growth via DVL1/Wnt/β-catenin signaling in breast cancer. Biochem Biophys Res Commun. 2018;505:302–8.

102. Di Leva G, Garofalo M, Croce CM. MicroRNAs in cancer. Annu Rev Pathol. 2014;9:287–314.

103. Zhou J, Li W, Guo J, Li G, Chen F, Zhou J. Downregulation of miR-329 promotes cell invasion by regulating BRD4 and predicts poor prognosis in hepatocellular carcinoma. Tumour Biol J Int Soc Oncodevelopmental Biol Med. 2016;37:3561–9.

104. Wang X, Lu X, Zhang T, Wen C, Shi M, Tang X, et al. mir-329 restricts tumor growth by targeting grb2 in pancreatic cancer. Oncotarget. 2016;7:21441–53.

105. Kang H, Kim C, Lee H, Rho JG, Seo J-W, Nam J-W, et al. Downregulation of microRNA-362-3p and microRNA-329 promotes tumor progression in human breast cancer. Cell Death Differ. 2016;23:484–95.

106. Li W, Liang J, Zhang Z, Lou H, Zhao L, Xu Y, et al. MicroRNA-329-3p targets MAPK1 to suppress cell proliferation, migration and invasion in cervical cancer. Oncol Rep. 2017;37:2743–50.

107. Zhao J, Tao Y, Zhou Y, Qin N, Chen C, Tian D, et al. MicroRNA-7: a promising new target in cancer therapy. Cancer Cell Int [Internet]. 2015 [cited 2020 Apr 23];15. Available from: https://www.ncbi.nlm.nih.gov/pmc/articles/PMC4625531/

108. Swarbrick A, Woods SL, Shaw A, Balakrishnan A, Phua Y, Nguyen A, et al. miR-380-5p represses p53 to control cellular survival and is associated with poor outcome in MYCN amplified neuroblastoma. Nat Med. 2010;16:1134–40.

109. Yang J-S, Maurin T, Robine N, Rasmussen KD, Jeffrey KL, Chandwani R, et al. Conserved vertebrate mir-451 provides a platform for Dicer-independent, Ago2-mediated microRNA biogenesis. Proc Natl Acad Sci U S A. 2010;107:15163–8.

110. Bailey MH, Tokheim C, Porta-Pardo E, Sengupta S, Bertrand D, Weerasinghe A, et al. Comprehensive Characterization of Cancer Driver Genes and Mutations. Cell. 2018;173:371–385.e18.

111. Fleming NI, Jorissen RN, Mouradov D, Christie M, Sakthianandeswaren A, Palmieri M, et al. SMAD2, SMAD3 and SMAD4 mutations in colorectal cancer. Cancer Res. 2013;73:725–35.

112. Korkut A, Zaidi S, Kanchi RS, Rao S, Gough NR, Schultz A, et al. A Pan-Cancer Analysis Reveals High-Frequency Genetic Alterations in Mediators of Signaling by the TGF-β Superfamily. Cell Syst. 2018;7:422–437.e7.

113. Massagué J, Wotton D. Transcriptional control by the TGF-beta/Smad signaling system. EMBO J. 2000;19:1745–54.

114. Massagué J. TGFβ signalling in context. Nat Rev Mol Cell Biol. 2012;13:616–30.

115. Nolan BE, Levenson E, Chen BY. Influential Mutations in the SMAD4 Trimer Complex Can Be Detected from Disruptions of Electrostatic Complementarity. J Comput Biol J Comput Mol Cell Biol. 2017;24:68–78.

116. Woodford-Richens KL, Rowan AJ, Gorman P, Halford S, Bicknell DC, Wasan HS, et al. SMAD4 mutations in colorectal cancer probably occur before chromosomal instability, but after divergence of the microsatellite instability pathway. Proc Natl Acad Sci U S A. 2001;98:9719–23.

117. Miyaki M, Iijima T, Konishi M, Sakai K, Ishii A, Yasuno M, et al. Higher frequency of Smad4 gene mutation in human colorectal cancer with distant metastasis. Oncogene. 1999;18:3098–103.

118. Koyama M, Ito M, Nagai H, Emi M, Moriyama Y. Inactivation of both alleles of the DPC4/SMAD4 gene in advanced colorectal cancers: identification of seven novel somatic mutations in tumors from Japanese patients. Mutat Res. 1999;406:71–7.

119. Salovaara R, Roth S, Loukola A, Launonen V, Sistonen P, Avizienyte E, et al. Frequent loss of SMAD4/DPC4 protein in colorectal cancers. Gut. 2002;51:56–9.

120. Blahna MT, Hata A. Smad-mediated regulation of microRNA biosynthesis. FEBS Lett. 2012;586:1906–12.

121. Tecalco-Cruz AC, Ríos-López DG, Vázquez-Victorio G, Rosales-Alvarez RE, Macías-Silva M. Transcriptional cofactors Ski and SnoN are major regulators of the TGF-β/Smad signaling pathway in health and disease. Signal Transduct Target Ther. 2018;3:15.

122. Wu JW, Hu M, Chai J, Seoane J, Huse M, Li C, et al. Crystal structure of a phosphorylated Smad2. Recognition of phosphoserine by the MH2 domain and insights on Smad function in TGF-beta signaling. Mol Cell. 2001;8:1277–89.

123. Souchelnytskyi S, Tamaki K, Engström U, Wernstedt C, ten Dijke P, Heldin CH. Phosphorylation of Ser465 and Ser467 in the C terminus of Smad2 mediates interaction with Smad4 and is required for transforming growth factor-beta signaling. J Biol Chem. 1997;272:28107–15.

124. Davis BN, Hilyard AC, Lagna G, Hata A. SMAD proteins control DROSHA-mediated microRNA maturation. Nature. 2008;454:56–61.

125. Warner DR, Bhattacherjee V, Yin X, Singh S, Mukhopadhyay P, Pisano MM, et al. Functional interaction between Smad, CREB binding protein, and p68 RNA helicase. Biochem Biophys Res Commun. 2004;324:70–6.

126. Wu MK, Vujanic GM, Fahiminiya S, Watanabe N, Thorner PS, O’Sullivan MJ, et al. Anaplastic sarcomas of the kidney are characterized by DICER1 mutations. Mod Pathol Off J U S Can Acad Pathol Inc. 2018;31:169–78.

127. Chen J, Wang Y, McMonechy MK, Anglesio MS, Yang W, Senz J, et al. Recurrent DICER1 hotspot mutations in endometrial tumours and their impact on microRNA biogenesis. J Pathol. 2015;237:215–25.

128. Morlando M, Dini Modigliani S, Torrelli G, Rosa A, Di Carlo V, Caffarelli E, et al. FUS stimulates microRNA biogenesis by facilitating co-transcriptional Drosha recruitment. EMBO J. 2012;31:4502–10.

129. Auyeung VC, Ulitsky I, McGeary SE, Bartel DP. Beyond secondary structure: primary-sequence determinants license pri-miRNA hairpins for processing. Cell. 2013;152:844–58.

130. Lee Y, Ahn C, Han J, Choi H, Kim J, Yim J, et al. The nuclear RNase III Drosha initiates microRNA processing. Nature. 2003;425:415–9.

131. Kim Y-K, Kim B, Kim VN. Re-evaluation of the roles of DROSHA, Export in 5, and DICER in microRNA biogenesis. Proc Natl Acad Sci U S A. 2016;113:E1881–1889.

132. Han J, Lee Y, Yeom K-H, Kim Y-K, Jin H, Kim VN. The Drosha-DGCR8 complex in primary microRNA processing. Genes Dev. 2004;18:3016–27.

133. Fukuda T, Yamagata K, Fujiyama S, Matsumoto T, Koshida I, Yoshimura K, et al. DEAD-box RNA helicase subunits of the Drosha complex are required for processing of rRNA and a subset of microRNAs. Nat Cell Biol. 2007;9:604–11.

134. Suzuki HI, Yamagata K, Sugimoto K, Iwamoto T, Kato S, Miyazono K. Modulation of microRNA processing by p53. Nature. 2009;460:529–33.

135. Gregory RI, Yan K-P, Amuthan G, Chendrimada T, Doratotaj B, Cooch N, et al. The Microprocessor complex mediates the genesis of microRNAs. Nature. 2004;432:235–40.

136. Fletcher CE, Godfrey JD, Shibakawa A, Bushell M, Bevan CL. A novel role for GSK3β as a modulator of Drosha microprocessor activity and MicroRNA biogenesis. Nucleic Acids Res. 2017;45:2809–28.

137. Tang X, Li M, Tucker L, Ramratnam B. Glycogen synthase kinase 3 beta (GSK3β) phosphorylates the RNAase III enzyme Drosha at S300 and S302. PloS One. 2011;6:e20391.

138. Lund E, Güttinger S, Calado A, Dahlberg JE, Kutay U. Nuclear export of microRNA precursors. Science. 2004;303:95–8.

139. Bohnsack MT, Czaplinski K, Görlich D. Exportin 5 is a RanGTP-dependent dsRNA-binding protein that mediates nuclear export of pre-miRNAs. RNA. 2004;10:185–91.

140. Wang J, Lee JE, Riemondy K, Yu Y, Marquez SM, Lai EC, et al. XPO5 promotes primary miRNA processing independently of RanGTP. Nat Commun. 2020;11:1845.

141. Hutvágner G, McLachlan J, Pasquinelli AE, Bálint E, Tuschl T, Zamore PD. A cellular function for the RNA-interference enzyme Dicer in the maturation of the let-7 small temporal RNA. Science. 2001;293:834–8.

142. Knight SW, Bass BL. A role for the RNase III enzyme DCR-1 in RNA interference and germ line development in Caenorhabditis elegans. Science. 2001;293:2269–71.

143. Song M-S, Rossi JJ. Molecular mechanisms of Dicer: endonuclease and enzymatic activity. Biochem J. 2017;474:1603–18.

144. Chendrimada TP, Gregory RI, Kumaraswamy E, Norman J, Cooch N, Nishikura K, et al. TRBP recruits the Dicer complex to Ago2 for microRNA processing and gene silencing. Nature. 2005;436:740–4.

145. Lee HY, Zhou K, Smith AM, Noland CL, Doudna JA. Differential roles of human Dicer-binding proteins TRBP and PACT in small RNA processing. Nucleic Acids Res. 2013;41:6568–76.

146. Lee Y, Hur I, Park S-Y, Kim Y-K, Suh MR, Kim VN. The role of PACT in the RNA silencing pathway. EMBO J. 2006;25:522–32.

147. Tomaselli S, Bonamassa B, Alisi A, Nobili V, Locatelli F, Gallo A. ADAR Enzyme and miRNA Story: A Nucleotide that Can Make the Difference. Int J Mol Sci. 2013;14:22796–816.

148. Trabucchi M, Briata P, Filipowicz W, Ramos A, Gherzi R, Rosenfeld MG. KSRP Promotes the Maturation of a Group of miRNA Precuresors. Adv Exp Med Biol. 2011;700:36–42.

149. Nowak JS, Choudhury NR, de Lima Alves F, Rappsilber J, Michlewski G. Lin28a regulates neuronal differentiation and controls miR-9 production. Nat Commun. 2014;5:3687.

150. Viswanathan SR, Daley GQ, Gregory RI. Selective blockade of microRNA processing by Lin28. Science. 2008;320:97–100.

151. Thornton JE, Du P, Jing L, Sjekloca L, Lin S, Grossi E, et al. Selective microRNA uridylation by Zcchc6 (TUT7) and Zcchc11 (TUT4). Nucleic Acids Res. 2014;42:11777–91.

152. Höck J, Meister G. The Argonaute protein family. Genome Biol. BioMed Central; 2008;9:210.

153. Meister G. Argonaute proteins: functional insights and emerging roles. Nat Rev Genet. 2013;14:447–59.

154. Rand TA, Petersen S, Du F, Wang X. Argonaute2 cleaves the anti-guide strand of siRNA during RISC activation. Cell. 2005;123:621–9.

155. Mourelatos Z, Dostie J, Paushkin S, Sharma A, Charroux B, Abel L, et al. miRNPs: a novel class of ribonucleoproteins containing numerous microRNAs. Genes Dev. 2002;16:720–8.

156. Meister G, Landthaler M, Peters L, Chen PY, Urlaub H, Lührmann R, et al. Identification of novel argonaute-associated proteins. Curr Biol CB. 2005;15:2149–55.

157. Kenny PJ, Zhou H, Kim M, Skariah G, Khetani RS, Drnevich J, et al. MOV10 and FMRP regulate AGO2 association with microRNA recognition elements. Cell Rep. 2014;9:1729–41.

158. Jin P, Zarnescu DC, Ceman S, Nakamoto M, Mowrey J, Jongens TA, et al. Biochemical and genetic interaction between the fragile X mental retardation protein and the microRNA pathway. Nat Neurosci. 2004;7:113–7.

159. Wan R-P, Zhou L-T, Yang H-X, Zhou Y-T, Ye S-H, Zhao Q-H, et al. Involvement of FMRP in Primary MicroRNA Processing via Enhancing Drosha Translation. Mol Neurobiol. 2017;54:2585–94.

160. Liu J, Rivas FV, Wohlschlegel J, Yates JR, Parker R, Hannon GJ. A role for the P-body component GW182 in microRNA function. Nat Cell Biol. 2005;7:1261–6.

161. Jakymiw A, Lian S, Eystathioy T, Li S, Satoh M, Hamel JC, et al. Disruption of GW bodies impairs mammalian RNA interference. Nat Cell Biol. 2005;7:1267–74.

